# The lymphatic system favours survival of a unique *T. brucei* population, and its invasion results in major host pathology

**DOI:** 10.1101/2023.04.10.536298

**Authors:** Henrique Machado, Mariana De Niz

**Affiliations:** Instituto de Medicina Molecular João Lobo Antunes, Faculdade de Medicina, Universidade de Lisboa, Lisboa, Portugal

## Abstract

*Trypanosoma brucei* colonize and multiply in the blood vasculature, as well as in various organs of the host’s body. Lymph nodes have been previously shown to harbour large numbers of parasites, and the lymphatic system has been proposed as a key site that allows *T. brucei* distribution through, and colonization of the mammalian body. However, visualization of host-pathogen interactions in the lymphatic system has never captured dynamic events with high spatial and temporal resolution throughout infection. In our work, we used a mixture of tools including intravital microscopy and *ex vivo* imaging to study *T. brucei* distribution in 20 sets of lymph nodes. We demonstrate that lymph node colonization by *T. brucei* is different across lymph node sets, with the most heavily colonized being the draining lymph nodes of main tissue reservoirs: the gonadal adipose tissue and pancreas. Moreover, we show that the lymphatic vasculature is a pivotal site for parasite dispersal, and altering this colonization by blocking LYVE-1 (i.e. surface receptor of lymphatic vessel endothelial cells) is detrimental for parasite survival. Additionally, parasites within the lymphatic vasculature have unique morphological and behavioural characteristics, different to those found in the blood, demonstrating a physical separation of environments across both types of vasculature. Finally, we demonstrate that the lymph nodes and the lymphatic vasculature undergoes significant alterations during *T. brucei* infection, resulting in oedema throughout the host’s body.

## Introduction

*Trypanosoma brucei* is an extracellular parasite transmitted by the bite of infected tsetse flies (*Glossina* spp.). These parasites are known to colonize and multiply in the blood vasculature, as well as in the extravascular space of several organs (reviewed in (Crilly and Mugnier, 2021; Silva Pereira et al., 2019)). Historical studies have shown that *T. brucei* parasites have a preference for the lymphatic system (Goodwin, 1970; Schuberg & Boeing, 1913). While we and others have identified large *T. brucei* chronic reservoirs in tissues including the g-WAT (De Niz et al., 2021; Trindade et al., 2021, 2016), pancreas (De Niz et al., 2021), brain (Coles et al., 2015; Myburgh et al., 2013), skin (Alfituri et al., 2020b; Capewell et al., 2016) and lungs (Mabille et al., 2022), the distribution as well as morphological and behavioural characteristics of *T. brucei* populations in the lymph nodes and the lymphatic vasculature, remains relatively understudied. Tanner *et al* reported in 1980 (Tanner et al., 1980) the enrichment of *T. brucei* in the lymph nodes of rats and proposed these sites as important sites for parasite replication and antigenic variation. While these observations were performed in fixed tissues and at one time point of infection, current technology allowed us to investigate parasites *in situ*, *in vivo* with high spatial and temporal resolution. In the present work we used intravital microscopy to investigate *T. brucei* in 20 sets of lymph nodes, and in the lymphatic vasculature of the whole mouse body. We asked a) whether parasite density varies, temporally and anatomically, throughout 20 days of infection across 20 lymph node sets and lymphatic vasculature studied in this work and how it compares to blood vasculature parasite density; b) whether the parasite population across lymph nodes differs to that in blood in terms of cell cycle, stumpy presence, morphology and behaviour; c) whether the 20 sets of lymph nodes and lymphatic vasculature are remodeled during infection and what the consequences are for systemic homeostasis and d) if exogenously modulating the vascular endothelium results in alterations to host-parasite interactions and survival.

## Results

### Parasite density is in lymph nodes is different to blood, and varies across lymph node groups

In this work we began by investigating *T. brucei* population density (expressed as number of parasites per cm^2^ of tissue) within 19 lymph node sets distributed in 5 groups: the head and neck (HN), forelimbs and hindlimbs (L), intra-thoracic (T), intra-abdominal (A), and pelvis (P) of C57BL/6J infected mice. The lymph nodes we studied include the mandibular, superficial parotid, cranial deep cervical, proper axillary, subiliac, sciatic, popliteal, tracheobronchial, gastric, jejunal, colic, pancreaticoduodenal, lateral iliac, medial iliac, external iliac, renal, lumbar, caudal mediastinal, and caudal mesenteric lymph nodes (schematic is shown in **Figure 1A**). We measured *T. brucei* density in these 19 lymph node sets over 20 days of infection, and compared it to the parasite density found in the blood vasculature. Using hierarchical clustering, we observed that the blood is clustered separately from any of the lymph node sets (**Figure 1B**). We then observed two large sub-clusters of lymph nodes: one which includes all lymph nodes of the head and neck, all lymph nodes of the limbs, 2 lymph nodes of the thorax (tracheobronchial and cranial mediastinal), and 2 lymph nodes of the pelvis (lumbar aortic and external iliac). The other large cluster includes all other pelvic and thoracic-abdominal lymph nodes as well as all abdominal lymph nodes. All statistical analyses of comparisons to blood, within and between groups of lymph nodes are included in **Table 1**.

**Figure 1.**
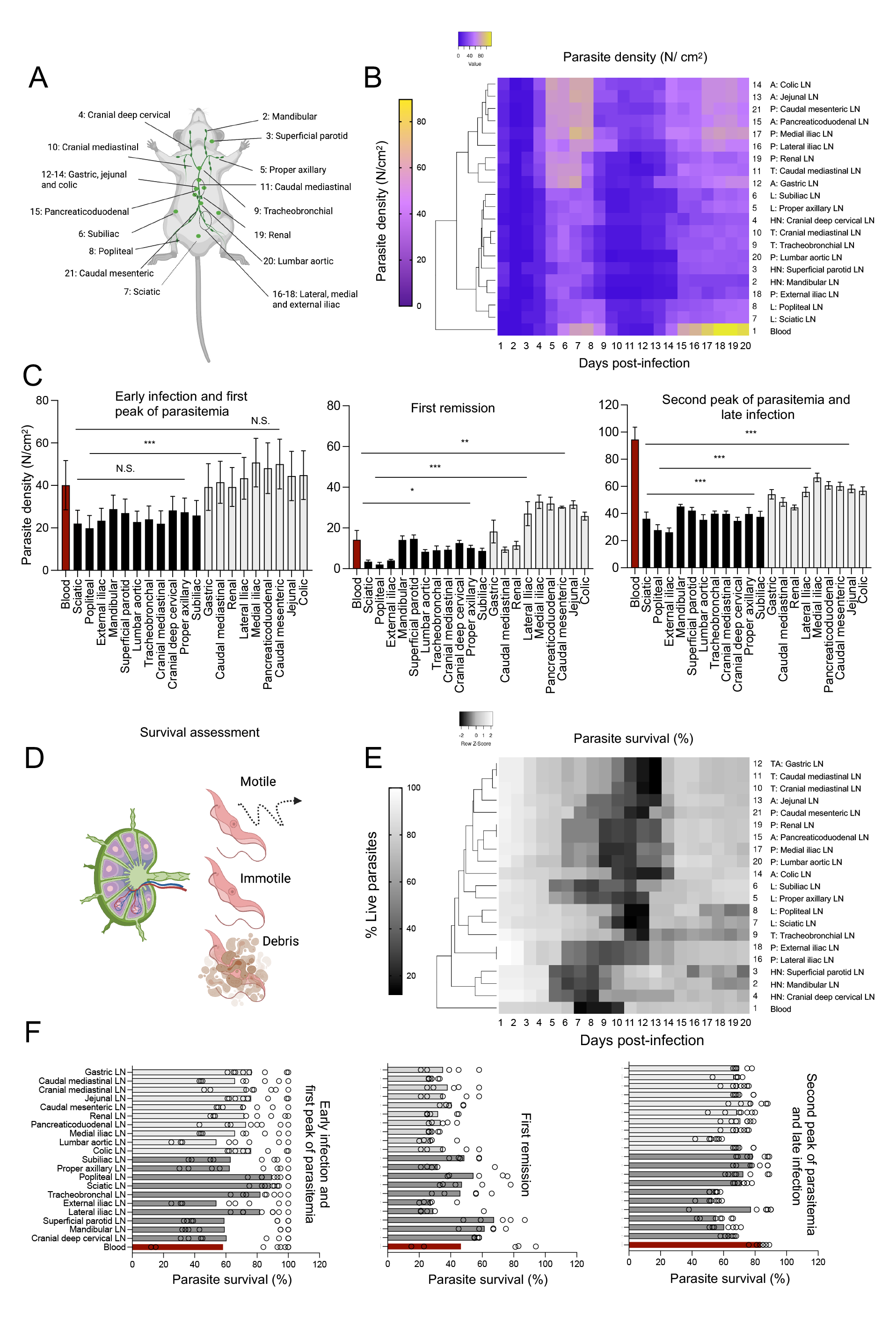
*T. brucei* density and survival across 20 lymph nodes and blood. **A)** Schematic representation of the anatomical localization of 20 lymph nodes distributed in the head and neck (HN, 2-4): mandibular, superficial parotid and cranial deep cervical; forelimbs and hindlimbs (grouped together as limbs (L, 5-8): proper axillary, subiliac, sciatic and popliteal; intra-thoracic (T: 9-11): tracheobronchal, cranial mediastinal and caudal mediastinal; intra-abdominal (A, 12-15): gastric, jejunal, colic and pancreaticoduodenal; and pelvic (P, 16-21): renal, lumbar aortic, lateral iliac, medial iliac, external iliac, and caudal mesenteric. For most subsequent figures, number 1 is given to blood (not marked in the schematic). **B-C)** Parasite density in 20 lymph nodes and blood. Parasite density was calculated by measuring the extravascular area in at least 100 separate fields of view in each lymph node, and quantifying the parasite number in this area. Quantities are expressed as parasite number (N) per cm^2^. Dendrogram shows the blood as a separate element, and 2 sub-clusters, one containing LNs 2-10, 18 and 20, and one containing LNs 11-17, 19 and 21. **C)** The average parasite density at 3 stages of infection (i.e. early infection and first peak of parasitemia (days 1-8); first remission (days 9-13); and second peak of parasitemia and late infection (days 14-20)) are shown for blood (red) and both lymph node sub-clusters (black bars and grey bars, respectively). One way ANOVA tests were performed to determine significant difference between clusters. Significance is shown as N.S. (non-significant), p ≤ 0.05 (*), p ≤ 0.01 (**) and p ≤ 0.001 (***). **D)** Graphical representation of features used to score survival in lymph nodes and blood: motility (whether productive motility leading to parasite displacement or not) was considered a main feature for the classification as live parasite. Completely immotile parasites (including lack of any indication of flagellar beating) that included signs of cell death (including loss of fluorescence and/or debris) were classified as dead. **E)** Parasite survival in 20 lymph nodes and blood. Parasite survival was calculated considering features mentioned in (D). Quantities are expressed as parasite percentage calculated in at least 100 fields of view. Dendrogram shows the blood as a separate element, with no clear LN sub-clusters. **F)** The average parasite survival at 3 stages of infection (i.e. early infection and first peak of parasitemia (days 1-8); first remission (days 9-13); and second peak of parasitemia and late infection (days 14-20)) are shown for blood (red) and two lymph node sub-clusters defined for (A-C) (dark grey and light grey bars, respectively). Circles represent the average parasite survival at each day post-infection.

Based on the 3 sub-clusters described above (blood and lymph node clusters 1 and 2), we went on to investigate relative parasite densities over time across lymph nodes (**Figure 1C, Figure S1, Table 2**). We explored parasite density in 3 time sets: early infection and first peak of parasitemia (days 1-8, left panel), first remission (days 9-13, middle panel) and second peak of parasitemia and late infection (days 14-20, right panel) (**Figure 1C**). On average, parasite burden during the early stage and first peak is 39.6, 24.6 and 44.6 parasites/cm^2^ in the blood and lymph nodes of cluster 1 and 2 respectively. The difference in parasite density between lymph node sub-clusters 1 and 2 at this stage, is statistically significant (p<0.01), but not the difference between either LN sub-cluster and blood (p > 0.05) (**Figure 1C, left panel; Table 2**). Analysis at higher temporal resolution showed that during this early stage, all lymph nodes are colonized faster than blood, and 24 hours after infection they have, on average 8.2-fold (cluster 1) and 25.5-fold (cluster 2) higher parasite densities than blood (**Figure S1A, Table 2**). Despite this faster initial colonization, parasite density in the blood rises quickly, reaching an average of 15 parasites/cm^2^ by day 3 post-infection, while lymph nodes in cluster 1 and 2 reach an average of 12.7 parasites/cm^2^ and 34 parasites/cm^2^ respectively. This represents a 192-fold increase relative to day 1 in the blood, a 30.3-fold increase in lymph nodes of cluster 1, and a 25.4-fold increase in lymph nodes of cluster 2 (**Figure S1B, Table 2**). Parasite density continues to rise until its first peak at days 7-8 post-infection in both the blood and lymph nodes of both clusters (**Figure 1B, Table 1**).

This initial phase is followed by remission, where the minimum parasite density (second to parasitemia at day 1) across all 20 days is reached in both blood, and lymph nodes of both clusters. The average parasite density during this stage is 14.5, 8.7 and 24.3 parasites/cm^2^ in the blood and lymph nodes of clusters 1 and 2 respectively. Differences between both LN sub-clusters and the blood, and between both LN sub-clusters are statistically significant (p <0.01) (**Figure 1C, middle panel; Table 2**). Finally, in the second peak of parasitemia, parasite density increases again in the blood and lymph node clusters 1 and 2 to 95, 36.7, and 56.2 parasites/cm^2^ respectively. Differences between both LN sub-clusters and the blood, and between both LN sub-clusters are statistically significant (p <0.001) (**Figure 1C, right panel; Table 2**). Although the cumulative average is highest during the 2^nd^ peak of parasitemia in LNs and blood, the absolute maximum parasite density across all days is reached in this 3^rd^ stage in the blood, but not in most lymph nodes (**Table 2, Figure S1C**), as most lymph nodes show the maximum parasite density around days 5 and 7 post-infection, yet the difference between the maximum density at the first and second peaks of parasitemia is non-significant. Altogether, the early stage of infection and the remission phase seem to favour lymph node invasion over blood, and throughout all 3 infection stages parasite density variations in the lymph nodes remain minimal (**Figure S1D**). The most parasite-enriched lymph nodes are proximal to the largest parasite reservoirs (e.g. pancreas and AT), and concentrate in the abdominal and pelvic regions of the mouse body.

Additional to total and relative parasite density, parasite survival in the lymph nodes varied greatly throughout infection. We scored viability using two criteria: motility and cell integrity. Loss of cell integrity included cell damage or signs of apoptosis, as well as debris (**Figure 1D**). Hierarchical clustering showed that the blood is a separate cluster from the lymph nodes (**Figure 1E**). Although large separate clusters of lymph nodes were not identified, a tendency towards lymph nodes within each anatomical group (i.e. HN, L, T, A, P) to show similar characteristics remained, and the grouping used for parasite density analysis was kept. Across all anatomical locations, parasite viability was maximal during early infection, and minimal around the first peak of parasitemia (prior to remission). Geometric means and individual points corresponding to each day are shown for three infection stages: early infection and first peak of parasitemia, remission, second peak of parasitemia and late infection. During early infection and first peak of parasitemia, the average parasite survival was 58.2, 67.8 and 73.1% in the blood and lymph node sub-clusters 1 and 2 respectively (**Figure 1F left panel, Table 3**). During remission this fell to 46.5, 46.0 and 33.1% in the blood and lymph node sub-clusters 1 and 2 respectively (**Figure 1F middle panel, Table 3**). This was followed by a rise in survival during the second peak of parasitemia and late infection, reaching 83.6, 64.2 and 71.6% survival in the blood and lymph node sub-clusters 1 and 2 respectively (**Figure 1F right panel, Table 3**). Although on average survival was highest in the blood, an absolute minimum of 12% survival was recorded in the blood, while the minimum survival in lymph node sub-clusters 1 and 2 was of 29.2 and 24.4% respectively (**Figure S1E**). Therefore, survival fluctuations were highest in the blood (as explored in **Table 3 and Figure S1F**). To avoid mortality-related confounding factors, only live parasites were considered for all further analysis. Altogether, based on these observations, we hypothesize that *T. brucei* successfully adapt to lymph nodes, and display high viability in these organs during infection.

### The mouse lymph nodes harbour a rapidly replicating parasite population, and are a less favourable environment for stumpy presence/formation

Having detected the different parasite densities and survival patterns of *T. brucei* across the 20 lymph node sets, we went on to ask whether the parasite population within the lymph nodes differs significantly from the bloodstream population and between lymph node sets. We began by analysing cell cycle progression among the parasites populating each lymph node set, by nucleus and kinetoplast quantification (**Figure 2A left panel**). Hierarchical clustering showed 2 major clusters, one of which includes the blood, all pelvic (P), all thoracic (T) and most abdominal (A) lymph nodes. The other cluster includes all head and neck (HN) lymph nodes. Limb lymph nodes are distributed across both major clusters (**Figure 2A, Table 4**). We found that on average, all lymph nodes have significantly lower percentages of parasites at 1K1N stage than the average parasite population in blood (p<0.01) (**Figure 2B, Table 4**). While the average percentage of parasites at 1K1N stage in the blood was 73.2%, it was 66.6, 66.2, 69.1, 67.9 and 68.1% in HN, L, T, A and P lymph node sets respectively. This difference between lymph nodes and blood was significant in all cases (**Table 4**). However, the difference within lymph node groups in HN, L, T, A and P was not significant. The average percentage of parasites at 2K1N stage in the blood was 18.6%, while it was 17.3, 17.9, 16, 16.9 and 18.3% in HN, L, T, A and P lymph node sets respectively. The difference between lymph nodes and blood was mostly not significant, while within most groups (all except L) the differences were significant. (**Figure 2C left panel, Table 4**). Finally, the average percentage of parasites at 2K2N stage in the blood was 8.4%, while it was 16.1, 15.8, 14.9, 15.2 and 13.7 % in HN, L, T, A and P lymph node sets respectively. This difference was significant for all lymph nodes compared to blood (p<0.001), but not significantly different when compared within lymph node groups, except for A lymph nodes (**Figure 2C right panel, Table 4**).

**Figure 2.**
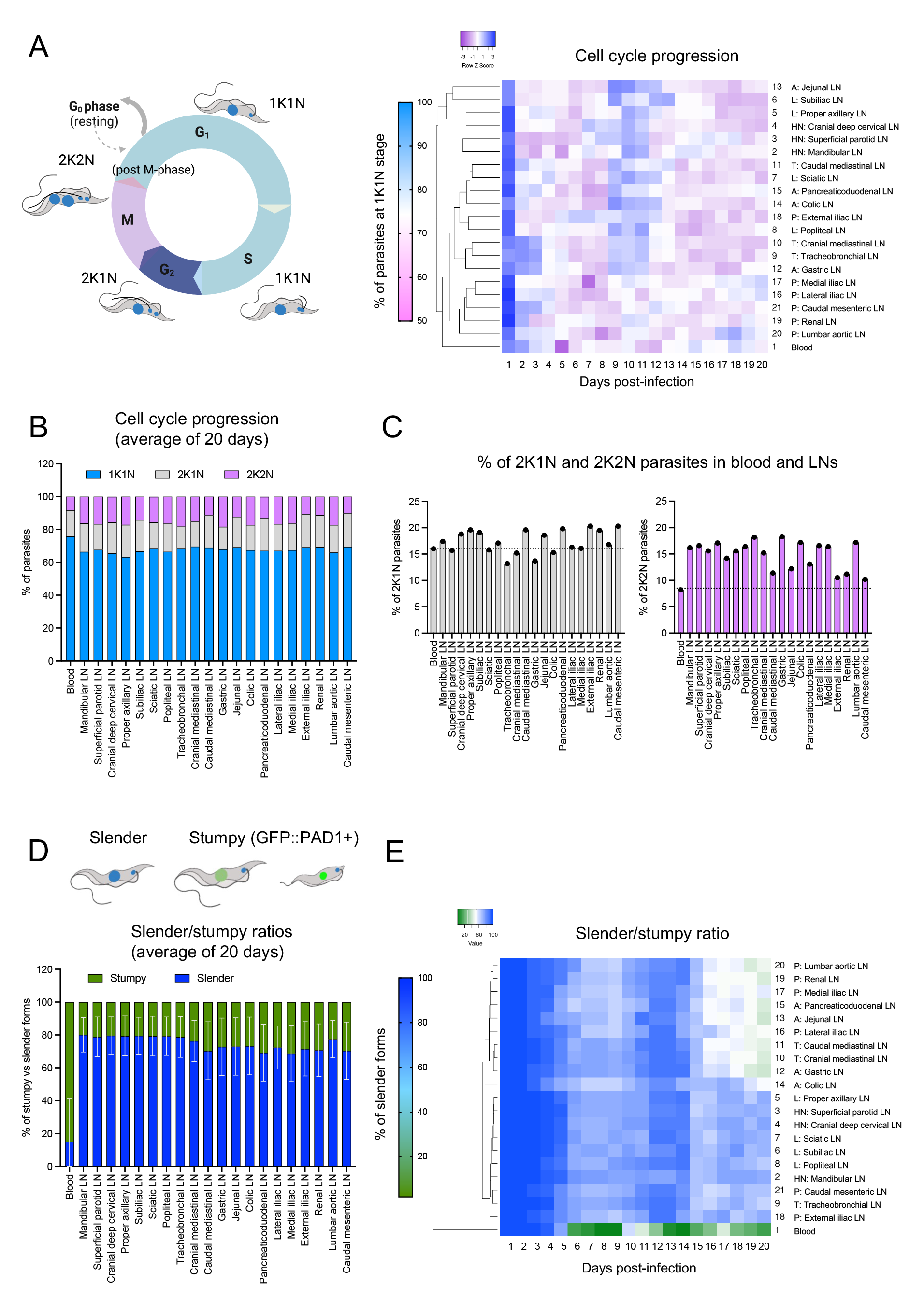
T. brucei cell cycle progression and stumpy/slender presence across 20 lymph nodes and blood. **A)** Schematic representation of *T. brucei* cell cycle, showing 1 kinetoplast and 1 nucleus (1K1N) in the G and S phase (shown in blue), 2K2N in the G2 phase (shown in dark blue), and 2K2N in M and the post-M phase (shown in purple). Parasite 1K1N, 2K1N and 2K2N percentages in 20 lymph nodes and blood were calculated in at least 100 separate fields of view. Dendrogram shows only the 1K1N population, and shows the blood as a separate element, with sub-clusters corresponding, to some extent, to anatomical location (i.e. HN, L, T, A, P). **B-C)** Relative percentages of each sub-population are shown for all organs measured. (C) shows the separate percentages of the 2K1N and 2K2N populations. The dotted line shows the blood average, which is not significantly different for the 2K1N population, but significantly different for the 2K2N population, as determined by student t-test for each LN relative to blood, and One-way ANOVA for LN groups relative to blood and to each other. **D)** Schematic representation of slender and stumpy *T. brucei* populations. For the purpose of this work, GFP-negative forms were defined as slender forms, while GFP-positive forms were defined as stumpy. Importantly, GFP-positive forms did not only include fully differentiated stumpy forms, however this was mostly relevant to the blood, as the majority of parasites in the LNs were GFP-negative. Slender (blue) and stumpy (green) percentages in 20 lymph nodes and blood were calculated in at least 100 separate fields of view. Graph shows the geometric mean of all 20 days of infection. Bars represent the geometric SD. **E)** Dendrogram for the slender/stumpy ratio over 20 days of infection shows the blood as a separate element, with two main sub-clusters, one including all HN and L lymph nodes, and one including all A and most P and T lymph nodes.

Given the significant difference in 1K1N and 2K2N forms in the blood compared to all lymph nodes, we hypothesized that differences might exist in the presence of stumpy and slender forms in the blood and lymph nodes. We went on to score the presence of stumpy and slender forms by the use of a PAD1:GFP reporter parasite line (**Figures 2D-2E**). Hierarchical clustering (**Figure 2E**) shows the blood as a separate cluster, and two main lymph node sub-clusters, one including all HN and L lymph nodes, and another including most of the T, A and P lymph nodes. We calculated the geometric mean of the percentage of slender forms in blood to be 15.01%, while it was 79.56 and 79.41% in lymph node cluster 1 (HN and L), and 75.1, 72.1, and 71.9% in lymph node cluster 2 (T,A and P) (**Figure 2D-2E, Table 5**). Temporally resolved quantifications showed significantly different patterns of slender/stumpy presence throughout infection. In the blood, there is a steep decrease in slender forms by the first peak of parasitemia (day 6-9), reaching the absolute minimum (i.e. stumpy maximum %) at days 8-9. During the first half of the infection, the average slender percentage is 54.6%, while in the second half it is 21.3% reaching a minimum of 2% (**Figure S2A, Table 5**). In lymph node cluster 1 (HN and L), slender presence remains significantly higher throughout infection, with an average of 85.7% during the first 10 days, 74.9% during the last 10 days, and a minimum of 63% at any point during the infection (which occurs between days 15-20). In lymph node cluster 2 (T, A and P), slender presence remains significantly higher than in blood throughout infection, with an average of 81.3% during the first 10 days, 67.4% during the last 10 days, and a minimum of 40% at any point during the infection (between days 16 and 20) (**Figure S2B**, **Table 5**). Together, these observations suggest that the lymph nodes are an important reservoir that favours parasite replication, which coincides with previous reports of higher *T. brucei* proliferation rates in rat lymph nodes (Tanner et al., 1980). Moreover, it suggests that the lymph node environment is less favourable for stumpy formation/presence than blood. This finding has key implications for our understanding of the role of extravascular reservoirs (the lymphatic system in particular) for *T. brucei* transmission.

### The mouse lymph nodes harbour a parasite population morphologically different to the one in blood

In addition to replication rate and stumpy/slender differences, we observed that parasites in the lymph nodes display an altered morphology compared to blood-circulating forms. Namely, in all lymph nodes analysed, parasites become progressively longer and wider than parasites conventionally found in blood (**Figures S3 and 3A-3B**). This is summarized in **Tables 6-8** and coincides with previous findings reported in the lymph nodes of infected rats (Tanner et al., 1980). Morphological changes were a continuous rather than a spontaneous event, with the parasite population displaying variable morphology throughout infection. Hierarchical clustering for width measurements showed the blood as a separate cluster to all lymph nodes, with no other significant clusters between the lymph node sets (**Figure S3A-S3B**). In the blood, the average parasite width throughout 20 days of infection was 2.13 µm. Width varied between a minimum of 1.4 µm at day 1 post-infection, and a maximum of 2.5 µm at days 18-19 post-infection. This represents a 78% increment in width. In the lymph nodes, the variation recorded was on average 46.6%, and ranged between 18.7% (Jejunal LN) and 75.9% (Cranial deep cervical). The average width in the blood parasite population (2.13 µm) was significantly lower than the width of parasites in all lymph nodes (**Table 6**), which was on average, 2.5 µm. In the lymph node groups previously defined by anatomical location (HN, L, T, A, P), the average parasite width was 2.51 µm (min: 1.88 µm, max 2.82 µm) in the HN group; 2.48 µm (min: 1.91 µm, max: 2.76 µm) in the L group; 2.47 µm (min: 1.99 µm, max: 2.77 µm) in the T group, 2.46 µm (min: 2.06 µm, max: 2.74 µm) in the A group and 2.45 µm (min: 1.80 µm, max: 2.8 µm) in the P group (**Figure S3B**). In all groups, changes in width were time-dependent, with the highest widths recorded at the peaks of parasitemia, and the smallest widths recorded either at the beginning of infection or during the remission phase (**Table 6**). Within-group differences were non-significant for L, T, A and P groups (p>0.05), but significant for the HN group (p = 0.04). Considering all lymph nodes across all time points, the differences in width were not significant (p = 0.1).

**Figure 3.**
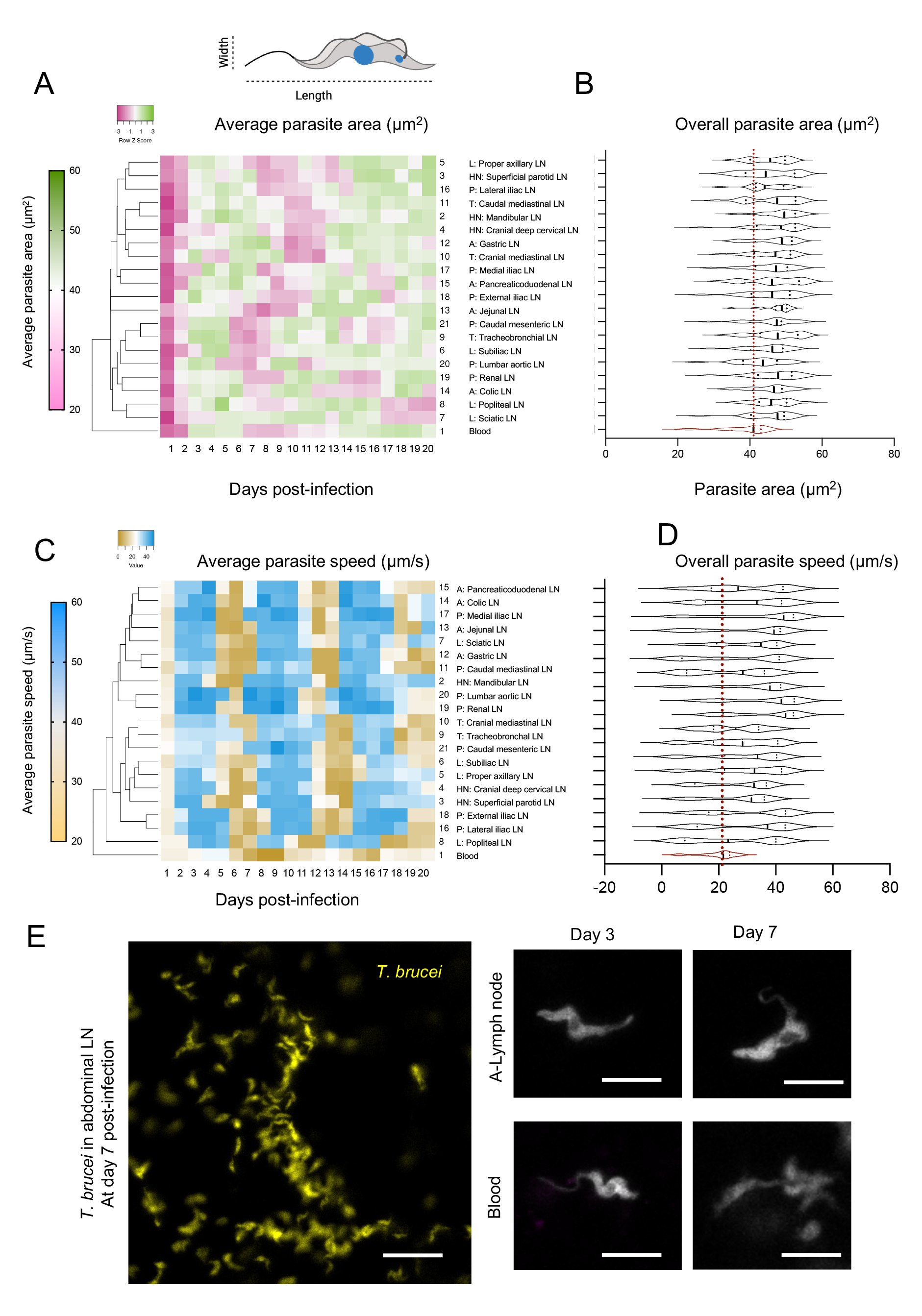
T. brucei morphology and behaviour across 20 lymph nodes and blood. **A)** Schematic representation of *T. brucei* length and width as used to calculate parasite area. Parasite length and width were calculated in at least 50 individual parasites in at least 20 fields of view in 20 lymph nodes and blood. Dendrogram shows the average parasite area at each day of infection. It shows the blood as a separate element, with no clear sub-cluster of lymph nodes. **B)** Violin plots showing the distribution of parasite areas for all organs analysed, during 20 days of infection. Blood is shown in red. **C)** Parasite speed was calculated in at least 50 individual parasites in at least 20 fields of view in 20 lymph nodes and blood. Dendrogram for the average parasite speed at each day of infection shows the blood as a separate element, with no major lymph node sub-clusters yet some degree of grouping by anatomical location (i.e. HN, L, T, A, P). **D)** Violin plots showing the distribution of parasite speed for all organs analysed, during 20 days of infection. Blood is shown in red. **E)** Representative images of parasite morphology in blood and LNs. Scale bar is 20 µm.

Similar to width, variations in parasite length also occurred throughout infection. Hierarchical clustering for length measurements showed the blood as a separate cluster to all lymph nodes, with no other significant clusters between the lymph node sets (**Figure S3C-S3D**). In the blood, the average parasite length throughout 20 days of infection was 17.7 µm. Length varied between a minimum of 15.76 µm at day 1 post-infection, and a maximum of 18.66 µm at day 17 post-infection (**Table 7**). This represents an 18.4% increment in length. In the lymph nodes, the variation recorded was on average 28.2%, and ranged between 21.5% (Cranial Deep Cervical LN) and 38.2% (Jejunal LN). The average length in the blood parasite population (17.7 µm) was significantly lower than the length of parasites in some lymph nodes (all in HN, popliteal (L), cranial mediastinal (T), gastric (A), jejunal (A), pancreaticoduodenal (A), external iliac (P), and caudal mesenteric (P) (**Table 7**), but not others. The average parasite length in all lymph nodes was 18.4 µm. In the lymph node groups previously defined by anatomical location (HN, L, T, A, P), the average, parasite length was 18.58 µm (min: 16.07 µm, max 20.02 µm) in the HN group; 18.18 µm (min: 15.75 µm, max: 20.02 µm) in the L group; 18.27 µm (min: 15.69 µm, max: 20.25 µm) in the T group; 18.83 µm (min: 15.9 µm, max: 20.68 µm) in the A group, and 18.25 µm (min: 15.49 µm, max: 19.96 µm) in the P group (**Figure S3D**). In all groups, changes in length were time-dependent, with the highest lengths recorded at variable times, but the smallest lengths recorded either at the beginning of infection or during the remission phase. Within-group differences were non-significant for HN, T and P groups (p>0.05), but significant for the L and A groups (p = 0.02 and 0.006 respectively). Considering all lymph nodes an all time points, the differences in length were significant (p = 0.01). While the average length was not significantly different between lymph nodes and blood in all cases, the maximum length reached by parasites in any lymph node (21.38 µm), was significantly different to the maximum reached in blood (18.66 µm). Equally, the difference between min. and max. length was highest in the lymph nodes (between 21.4 and 38.2% increase in length) compared to blood (18.4%) (**Table 7**).

Considering length and width parameters, we went on to determine whether the parasite area (defined as a proxy) was similar in blood and lymph nodes (**Figure 3A-B, Table 8**). Hierarchical clustering for area measurements showed the blood as a separate cluster to all lymph nodes, with no other significant clusters between the lymph node sets (**Figure 3A**). In the blood, the average parasite area throughout 20 days of infection was 37.5 µm^2^. Parasite area ranged between a minimum of 21.9 µm^2^ at day 1 post-infection, and a maximum of 45.49 µm^2^ at day 18 post-infection. This represents a doubling in area (>100% increment). In the lymph nodes, the area variation recorded was on average 74.9%, and ranged between 47.6% (Proper axillary LN) and 109.2% (Cranial Deep Cervical LN). The average area in the blood parasite population (37.54 µm^2^) was significantly lower than the area of parasites in all lymph nodes (**Table 8**). The average parasite area in all lymph nodes was 45.9 µm^2^. In the lymph node groups previously defined by anatomical location (HN, L, T, A, P), the average parasite area was 46.71 µm^2^ (min: 31.66 µm^2^, max 55.29 µm^2^) in the HN group; 45.19 µm^2^ (min: 31.51 µm^2^, max: 53.2 µm^2^) in the L group; 46.16 µm^2^ (min: 31.99 µm^2^, max: 54.35 µm^2^) in the T group, 46.46 µm^2^ (min: 34.07 µm^2^, max: 53.82 µm^2^) in the A group, and 44.79 µm^2^ (min: 28.37 µm^2^, max: 53.52 µm^2^) in the P group (**Figure 3B, Table 8**). Consistent with our separate findings for width and length, in all groups, changes in area were time-dependent, with the highest areas recorded during parasitemia peaks, and the smallest areas recorded mostly at the beginning of infection. Within-group differences were non-significant between lymph node groups (p>0.05). Considering all lymph nodes at all time points, the differences in parasite area were non-significant (p = 0.13). The maximum area reached by parasites in any lymph node (51.6 µm^2^ to 56.5 µm^2^), was significantly different to the one reached in blood (45.49 µm^2^) (**Table 8**). Notably, for all cases, parasite measurements were done in morphological slender forms. Altogether, we conclude that the parasite population in the lymph nodes is significantly different to the one found in blood, with parasites in lymph nodes having larger areas than those in blood.

### The mouse lymph nodes harbour a fast-moving parasite population behaviourally different to the one in blood

In addition to morphological changes, we went on to measure speed of motion (displacement as a function of time) in blood and lymph nodes (**Figure 3C-D, Figure S4, Table 9**). Hierarchical clustering for speed showed the blood as a separate cluster to all lymph nodes, without clear major sub-clusters arising for the LNs (**Figure 3C**). The geometric mean of parasite speed in blood throughout 20 days of infection was 14.74 µm/s. Parasite speed ranged between a minimum of 2.13 µm/s at the beginning of the remission phase, and a maximum of 27.46 µm/s at day 4 post-infection. This represents a 27-fold change in speed range. In the lymph nodes, parasites showed drastic changes in speed but the minimum speed was significantly higher than in blood. Considering all lymph nodes, the average speed was of 24.78 µm/s. The average difference between the minimum and maximum registered speed was a 7.26- fold change, with the highest range registered for the HN lymph nodes (10.56-fold change), and the lowest for T lymph nodes (5.15-fold change). The average parasite speed in the blood population (17.69 µm/s) was significantly lower than the speed of parasites in all lymph nodes (**Table 9**). In the lymph node groups previously defined by anatomical location (HN, L, T, A, P), the geometric mean of parasite speed was 24.76 µm/s (min: 4.07 µm/s, max: 42.95 µm/s) in the HN group; 23.36 µm/s (min: 6.89 µm/s, max: 42.13 µm/s) in the L group; 27.39 µm/s (min: 10.67 µm/s, max: 46.61 µm/s) in the T group; 21.09 µm/s (min: 5.91 µm/s, max: 41.24 µm/s) in the A group, and 26.9 µm/s (min: 7.29 µm/s, max: 47.3 µm/s) in the P group (**Figure 3D**). Consistent with our findings for differences in morphology, in all groups, changes in speed were infection time-dependent with drastic variations occurring within a short period of time during both parasitemia waves (**Figure S4A-D, Table 9**). Within-group differences were non-significant between most lymph node groups (p>0.05) except the T lymph nodes (p = 0.02). Considering all lymph nodes an all time points, the differences in parasite speed were significant (p = 0.04). Altogether, the maximum speed reached by parasites in any lymph node (50.92 µm/s), was significantly different to the one reached in blood (27.46 µm/s) (**Figure S4C-D, Table 9**). Representative images of parasites in blood and lymph nodes are shown in **Figure 3E**. Altogether, our data shows that in addition to morphological differences, the parasite population in the lymph nodes is significantly different to the one found in blood, with parasites displaying faster swimming in lymph nodes.

Finally, we considered all previously explored criteria (i.e. parasite density, parasite survival, cell cycle progression, stumpy/slender proportions, length, width, area, and speed) to perform a global analysis of differences and similarities between parasites in lymph nodes and blood, and between different lymph node sets (**Figure 4**). **Figure 4A** shows a distance matrix, which allowed us to explore whether altogether a) the lymph node parasite population was different to the one in the blood, and b) whether any sub-clustering existed within the 19 lymph node sets explored. The distance matrix showed the highest difference for blood (12.79). This is consistent with the most of the analyses done for each independent criteria. We then performed feature correlation and selection methods to determine which of the features analysed were most relevant for predicting parasite location (i.e. between blood and lymph nodes, as well as within lymph node sets. A correlation matrix (**Figure 4B, Table 10**) shows high correlation scores (>0.5) between parasite density, survival and cell cycle progression, as well as between parasite length, width, area and speed. Slender % has high negative correlation scores (<-0.4) relative to density, survival and cell cycle progression.

**Figure 4.**
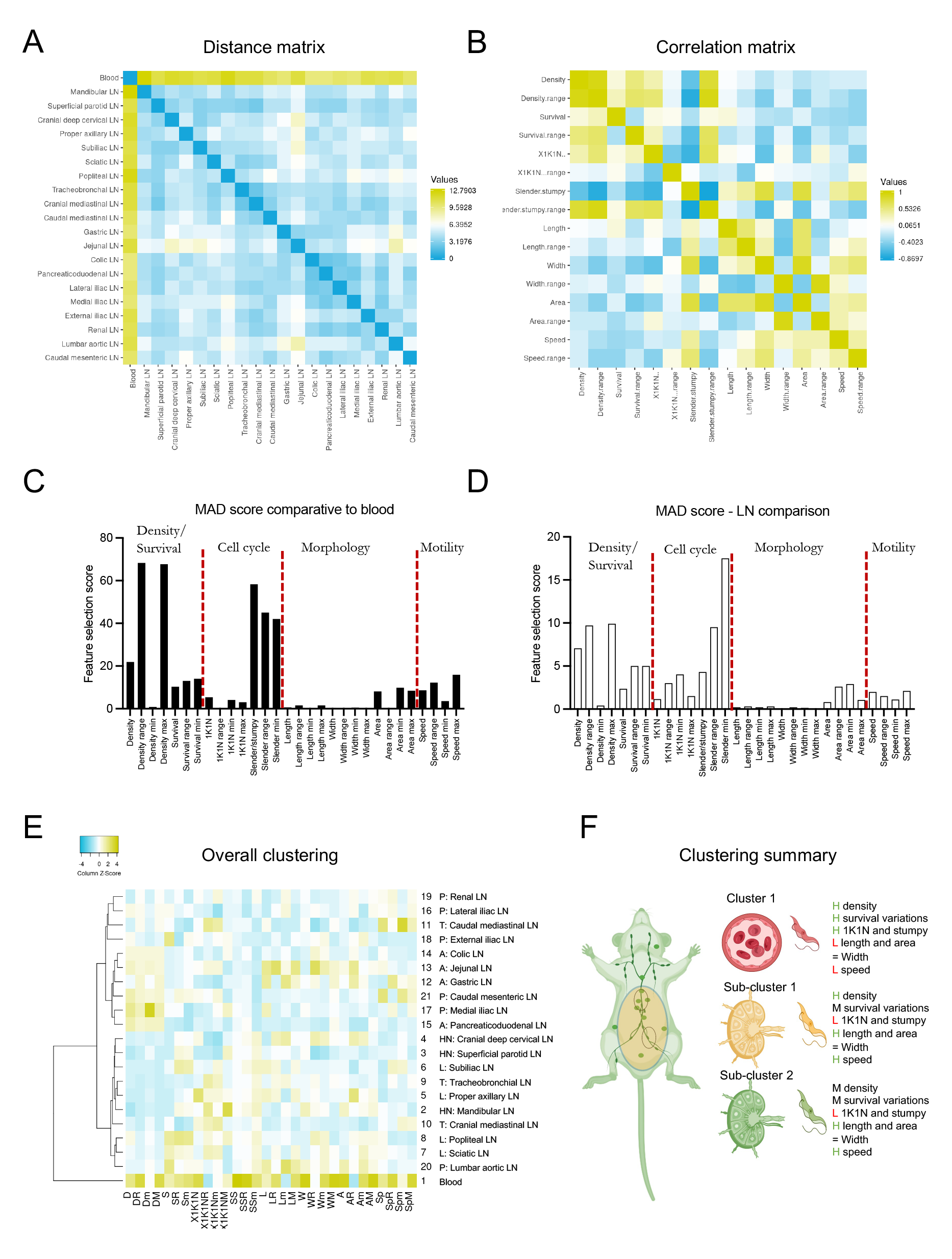
*T. brucei* blood population is significantly different to LN population, with the most defining selection factors being parasite density and slender/stumpy ratios. Considering parasite density, survival, cell cycle progression, slender/stumpy ratios, length, width, area and speed, including average, range, minimum and maximum values, various factor selection features were calculated. **A)** Distance matrix based on all aforementioned factors shows that the largest separation (12.8) exists for blood relative to any lymph node, while the distance between any lymph node is lower than 6. **B)** A correlation matrix shows high positive correlation (>0.5) between density, survival, and cell cycle progression, as well as between length, width, area and speed. **C)** Mean Absolute Deviation was used for factor selection calculation between blood and LNs. The highest score was reached by parasite density range and maximum, followed by slender/stumpy average, range and minimum. **D)** Mean Absolute Deviation was used for factor selection calculation between LNs. The highest score was reached by slender/stumpy minimum and range, followed by density range and density maximum. **E)** Considering all aforementioned features we investigated whether lymph node clustering existed. We confirmed the existence of 3 sub-clusters: one defined by the blood, one defined by all HN and L lymph nodes (as well as a P and 2 T lymph nodes), and one defined by all A and remaining P LNs). **F)** Schematic representation of overall findings regarding parasite populations in the LNs and blood.

For feature selection, we performed a MAD (median absolute deviation) analysis, which showed that key the most important features for distinguishing blood from lymph node *T. brucei* populations (**Figure 4C, Table 11**), include features relative to parasite density (specifically the density range (difference between maximum and minimum) and the density maxima), with MAD scores of 68.3 and 67.7 respectively. These were followed by features related to slender/stumpy ratios including the average, the minimum slender percentage registered, and the range), with MAD scores of 58.3, 42 and 45, respectively. Amongst the remaining features, speed range and maxima had the highest MAD scores (12.3 and 15.9 respectively). Considering only inter-lymph node comparisons, the same features as those previously discussed (i.e. parasite density and slender/stumpy ratios) had the highest MAD scores, with the slender/stumpy minima having the highest score (17.5). Overall, MAD scores for inter-LN differentiation were on average 4-fold lower than those recorded for the blood/LN differentiation (**Figure 4D, Table 11**).

While previously we investigated each feature independently to determine whether any clusters existed among LN sets and blood, we went on to investigate clustering based on a composite comparison. Hierarchical clustering considering all criteria (i.e. parasite density, parasite survival, cell cycle progression, stumpy/slender proportions, length, width, area, and speed) confirmed the blood as a separate cluster, and showed two large lymph node sub-clusters, one including all HN and L as well as most T lymph nodes, and one including all A and all P lymph nodes (**Figure 4E-4F**). This is consistent with most findings recorded for individual criteria, and highlights an important difference between lymph node groups based on their anatomical location, and the organ reservoirs they neighbour.

### Infection-induced structural remodelling of the lymphatic system correlates with accumulation of parasitised free fluid

In addition to our investigation on lymph node colonization and remodelling upon *T. brucei* infection, we also observed that the lymph nodes are significantly remodelled throughout infection (**Figure 5A-5B, Table 12**). Lymph node diameters were measured in uninfected mice, and at days 5, 10, 15 and 20 post-infection. Hierarchical clustering identified 3 major sub-clusters (**Figure 5A**). Although there was no clear sub-division coinciding fully with anatomical location, there was still a tendency for some lymph nodes to cluster together. One sub-cluster included most HN and all L lymph nodes, another included all A and most P lymph nodes, and one included most T and the remaining P lymph nodes. The average uninfected LN diameter was 1.6 mm, with HN, L and T lymph nodes being smaller (1.3, 1.4, 1.6 mm respectively) than A and P lymph nodes (1.9, 1.8 mm respectively). Most lymph nodes showed a significant increase in size by day 5 post-infection (p < 0.001) to an average diameter of 1.8 mm; to 2.4 mm by day 10 post-infection (p < 0.001); to 2.9 mm by day 15 post-infection (p < 0.001), and to 3.4 mm by day 20 post-infection (p < 0.001). This represented a 2.2-fold increase in size by day 20 post-infection, on average, with the greatest increase observed in T, A and P lymph nodes (**Figure 5B, Table 12**).

**Figure 5.**
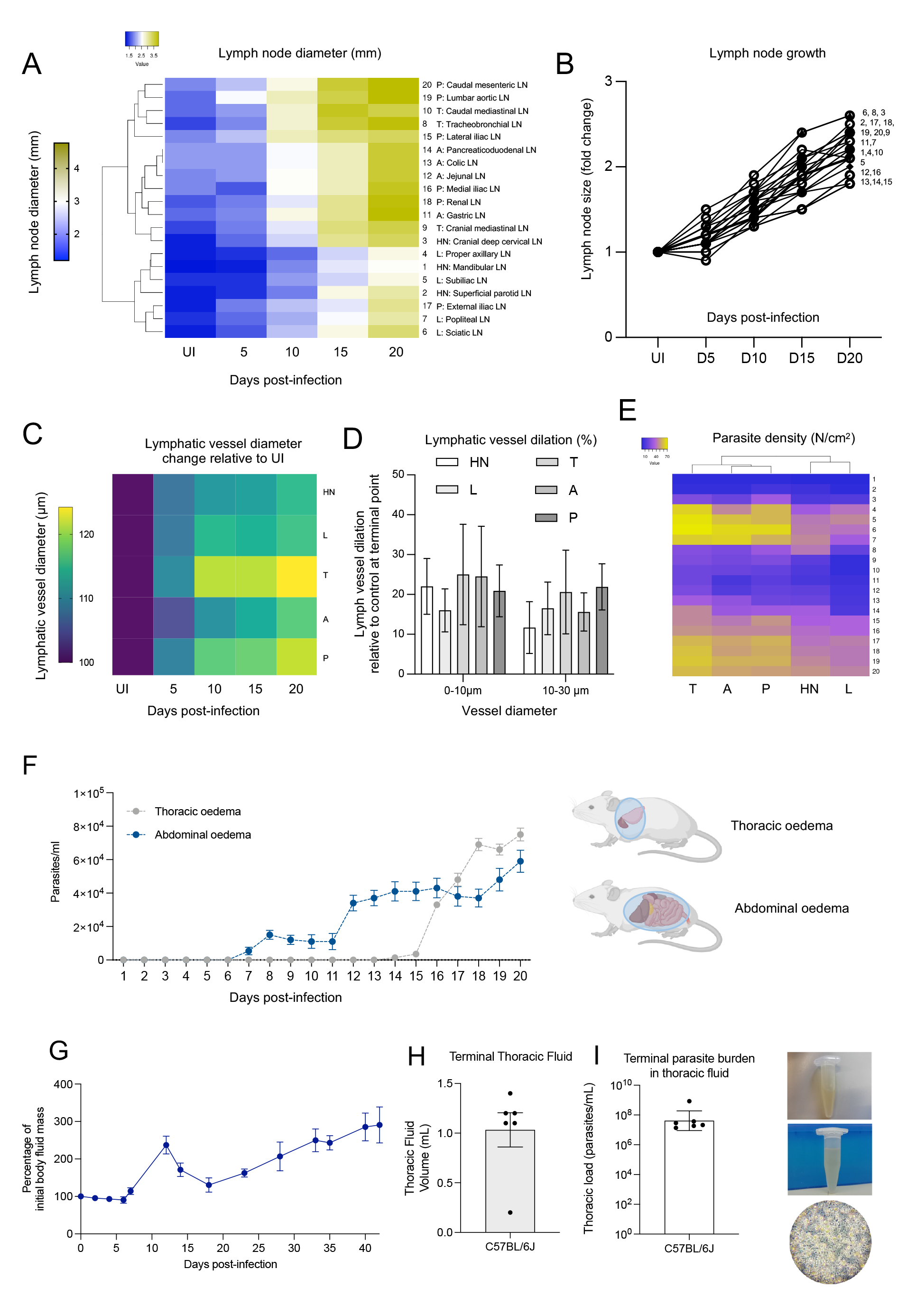
T. brucei infection results in significant lymphatic system remodeling and deleterious host pathology. **A)** Lymph nodes diameters were measured at 4 different times of infection, namely at days 5, 10, 15 and 20, as well as in uninfected mice. Hierarchical clustering showed 3 clusters with slightly different patterns of lymph node remodeling: most HN and L lymph nodes underwent significantly lower and more gradual enlargement than LNs in the T, A, and P areas. **B)** Analysis on the fold change in size by day 20 relative to uninfected mice shows that most lymph nodes increase their size by between 1.8 and 2.6-fold. Moreover, LN s continue to increase in size throughout infection. **C)** The lymphatic vasculature diameter was measured in one representative vasculature for each anatomical location (HN, L, T, A, P). An increase of between 15 and 24% in diameter was recorded for all representative lymphatic vessels, with the T and P lymphatic vasculature showing the greatest dilation. **D)** Further exploring whether dilation occurred equally in vessels of different diameter, although we observed a tendency towards smaller vessels (0-10 µm) to undergo greater dilation than larger vessels (10-30 µm), this difference was not statistically significant. Error bars shown represent SD. **E)** Parasite density in the lymphatic vasculature was calculated in HN, L, T, A, P locations, with T, A, P (the most highly dilated vasculature) showing highest parasite density. **F)** Parasitemia (expressed as parasites/ml) was calculated in the free fluid found in the thorax (grey lines) and abdomen (blue lines) of mice at each point of intravital surgery. While parasitemia in the free abdominal fluid gradually increased from day 6, as infection progresses, parasitemia in the thorax showed a sharp increase from day 15 of infection. By day 20 post-infection, the free abdominal fluid had a parasitemia of 5.9 x 10^4^ parasites/ml, while the free thoracic fluid reached 7.5 x 10^4^ parasites/ml. Dots represent average parasitemia. Error bars shown represent SD. **G)** Total Free fluid mass was measured by nuclear magnetic resonance throughout infection and then normalized to baseline. Free fluid mass increased by 2.9-fold by day 42 post-infection. (n=5 mice). **H)** Parasite burden of thoracic fluid and (**I**) volume isolated from moribund mice (33-51 days post-infection, n=6 mice).

We then went on to investigate whether the lymphatic vasculature also experienced anatomical changes throughout infection. We focused on vessel dilation, and selected vasculature of various diameters per anatomical location, namely, the mandibular LN (HN), the popliteal LN (L) the tracheobronchial LN (T), the pancreaticoduodenal LN (A), and the external iliac LN (P). For lymphatic vessel identification, we injected mice with A647-conjugated LYVE-1 antibody. Irrespective of anatomical location, we identified a significant lymphatic vessel dilation by day 5 post-infection, with significant increases every 5 days of infection (**Figure 5C, Table 13**). At the terminal point of infection (day 20), we identified that lymphatic vessels of medium diameter (0-10 µm) were enlarged by 22 % (HN), 16% (L), 25% (T), 25% (A) and 21% (P) relative to uninfected mice. While larger lymphatic vessels (10-30 µm) also showed significant enlargement, it was not as prominent as the one found in smaller vessels. Larger lymphatic vessels showed a 11.7% (HN), 16.5% (L), 20.5% (T), 15.6% (A), and 21.9% (P). Altogether, vessels linked to the T, A and P lymph nodes showed the greatest modifications (**Figure 5D, Table 13**). Although we explored whether vessel enlargement and parasite density (in the lymphatic vasculature) (**Figure 5E**) were correlated, we found an overall R^2^ value of 0.51, suggesting weak correlation.

Given this result suggesting lymph node and lymphatic vessel pathology, as well as our constant findings during intravital surgeries, of the presence of liquid consistent with oedema in the thoracic, and abdominal-pelvic cavities, we went on to analyse the isolated fluid from these compartments (**Figure 5F, Table 14**). Significant free fluid was detectable in the abdomen from day 7 post-infection (i.e. around the first peak of parasitemia), while it was only detectable in significant amounts in the thorax from day 14 post-infection. Upon analysis of the fluid for presence of parasites, we found 5.4 x 10^3^ parasites/ml and 1.4 x 10^3^ parasites/ml in the abdomen and thorax respectively. This increased 10-fold and 50-fold by day 20 post-infection in the abdomen and thorax, respectively. This is consistent with a gradual increase of parasites in the abdomen (1.3-fold daily), but a drastic increase in free fluid and parasite presence in the thorax (2.8-fold daily). Upon investigating free fluid consistent with lymphatic system pathology until day 42 post-infection, we observed fluctuations between −10% and +62% within the first 20 days of infection, relative to free fluid in uninfected mice. This drastically increased during the next 22 days culminating in +190% at day 42 post-infection relative to uninfected mice (**Figure 5G, Table 14**). At this point of infection, a large amount of free fluid was extracted from the thorax of moribund infected mice (i.e. between 0.2 and 1.4 ml), and this fluid was found to be heavily parasitized, with between 1.37 x 10^7^ and 8.55 x 10^8^ parasites/ml (**Figures 5H-I, Table 14**). Together, these findings show that a *T. brucei* infection leads to significant structural remodelling of the lymphatic system, which correlates with the accumulation of potentially lethal amounts of parasitised oedema.

### Blocking the lymphatic vascular receptor LYVE-1 is detrimental for parasite survival

Based on our findings regarding host pathology and parasite survival related to the colonization of the lymphatic system, we went on to explore the effect of blocking the lymphatic vascular endothelial hyaluronan receptor 1 (LYVE-1), on parasite spread and survival, and on host peripheral parasitemia and survival (**Figure 6, Table 15**). LYVE-1 is a type I integral membrane glycoprotein that contains hyaladherin and can bind hyaluronic acid, and is present as a cell surface receptor on lymphatic endothelial cells (Adamczyk et al., 2016; Jackson, 2019) (**Figure 6A**). Given the large presence of *T. brucei* parasites in the lymphatic system, we used blocking/neutralizing antibodies to investigate the effect of the LYVE-1 receptor (or lack thereof) on parasite distribution and survival (**Figure 6A-6D**). For this purpose, we injected α-LYVE-1 antibody every two days to ensure the receptor was neutralized until day 10 post-infection. We observed that upon loss of available LYVE-1 receptor, parasite density displays a different dynamic to isotype-treated or untreated mice, with α-LYVE-1-treated mice displaying a sharp increase in parasitemia which was 2-fold higher than isotype-treated or untreated mice until day 7 post-infection. This was followed by a sharp loss in viability from days 8-10 post-infection (**Figures 6B-6C**). While the geometric mean of the global parasite density was not significantly different (**Table 15**), the maxima (reached between days 4 and 7 post-infection) (**Figure 6D**), and minima (reached between days 8 and 10) were significantly different in all anatomical locations evaluated (i.e. HN, L, T, A, P) (p<0.05) (**Table 15**). The maximum parasite density in the lymphatic vasculature reached in untreated, isotype-treated and α-LYVE-1-treated mice was of 59, 58.4, and 82.3 parasites/cm^2^, respectively. The minimum parasite density (post-peak of parasitemia) in the lymphatic vasculature reached in untreated, isotype-treated and α- LYVE-1-treated mice was of 13.4, 12.9, and 4.3 parasites/cm^2^, respectively.

**Figure 6.**
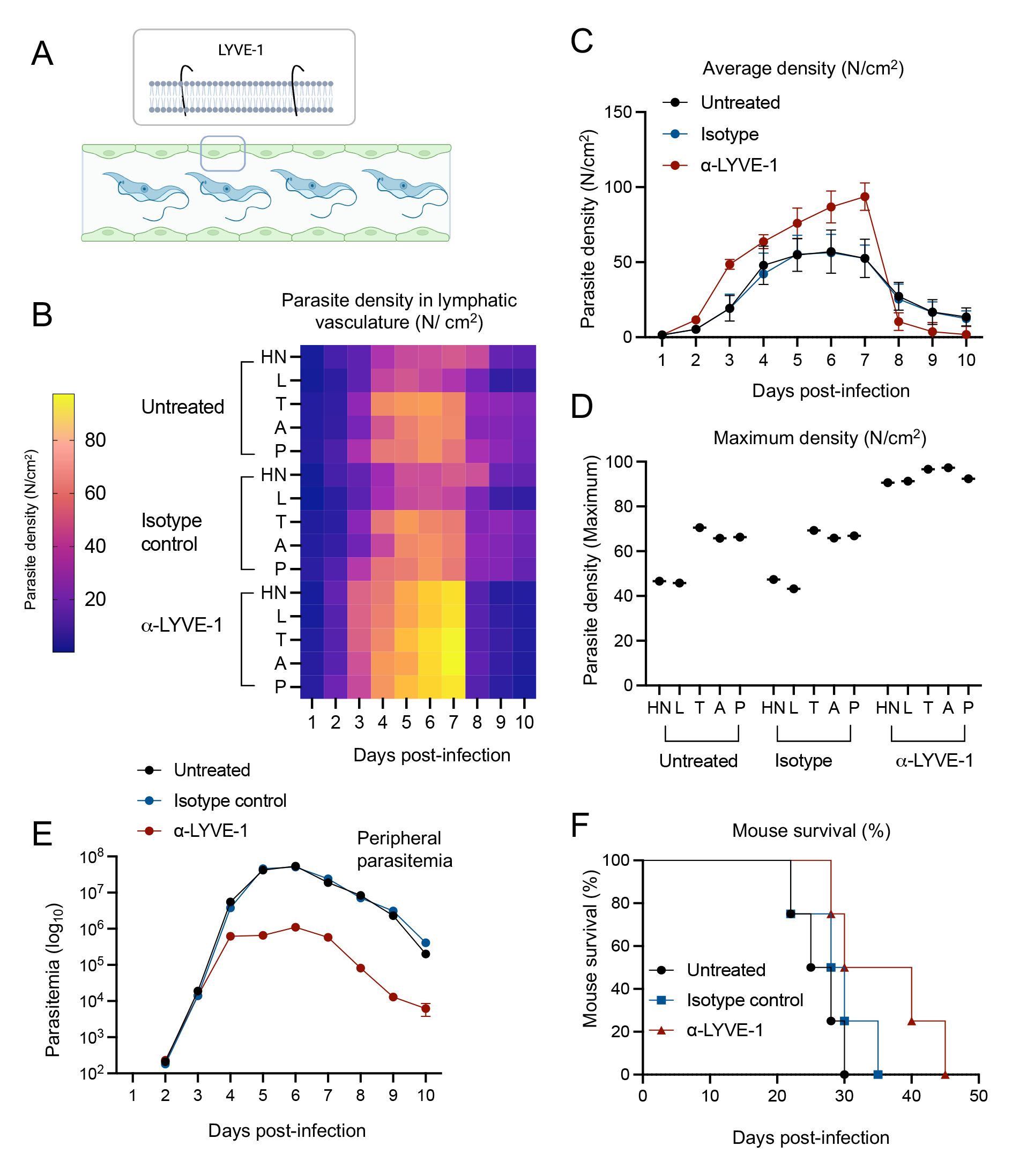
The lymphatic vascular endothelial receptor (LYVE-1) plays an important role in parasite colonization of the lymphatic system and blood. **A)** Schematic representation of LYVE-1, a type I integral membrane glycoprotein present in the lymphatic vascular endothelium. **B-C)** Parasite density (expressed as parasite number per area) was measured in the lymphatic vasculature of untreated mice, and mice treated with isotype control, and anti-LYVE-1 antibody to block the receptor. Parasite density was measured in lymphatic vasculature of the HN, L, T, A, P anatomical regions. Heat map shows that anti-LYVE-1 treated mice reach higher parasite densities between days 4 and 7 post-infection in all anatomical locations, and lowest parasite densities between days 8 and 10. **C)** Numerical representation of the heatmap shown in (B). Untreated values are shown in black, isotype control in blue, and anti-LYVE-1 in red. Parasite density in the lymphatic vasculature of anti-LYVE-1-treated mice experiences drastic changes, with the highest maxima and the lowest minima compared to both control groups. **D)** Parasite density maxima were significantly different between anti-LYVE-1-treated mice (93.6 parasites/cm^2^) and mice in both control groups (59 and 58.5 parasites/cm^2^). **E)** Blood parasitemia was significantly impacted by treatment with anti-LYVE-1 blocking antibodies. Control groups (untreated and isotype control-treated) reached an average parasitemia of 1.46 and 1.5 x 10^7^ parasites/ml throughout 10 days of infection, with maxima of 5.4 x 10^7^ and 5.1×10^7^ respectively. Conversely, anti-LYVE-1-treated mice reached an average parasitemia of 3.4 x 10^5^ parasites/ml, with a maximum of 1.1×10^6^ parasites/ml. Dots show the average parasitemia at each day of infection. Error bars represent SD. **F)** Mouse survival was significantly impacted by anti-LYVE-1-treatment, with control groups surviving between 20 and 35 days post infection, and anti-LYVE-1-treated groups surviving between 30 and 45 days post-infection.

Based on this result in the lymphatic vasculature, we went on to investigate whether peripheral parasitemia (in the blood vasculature) was affected by α-LYVE-1-treatment (**Figure 6E, Table 16**). We found that α-LYVE-1- treated mice have an overall 42.7-fold lower parasitemia than untreated mice, and 44-fold lower than isotype-control-treated mice (**Table 16**). Untreated and isotype-control-treated mice reached the peak of parasitemia at day 6 post-infection (5.4×10^7^ and 5.1×10^7^ parasites/ml respectively). At the same day post-infection, parasitemia in α-LYVE-1- treated mice was 1.1×10^6^ parasites/ml. Although we stopped administration of the blocking antibody at day 10 post-infection, we went on to evaluate the effect of this treatment on mouse survival. We observed that untreated mice and isotype-control-treated mice lived between 20 and 35 days post-infection, while α-LYVE-1-treated mice lived between 30 and 50 days (**Figure 6F**). This difference was significant (p = 0.03), and suggests an important interplay between the lymphatic system and general homeostasis during *T. brucei* infection.

## Discussion

The interplay between *T. brucei* and host lymph nodes has been an important focus of interest in *Trypanosoma* research since very soon after the discovery of the parasite by Sir David Bruce in 1894. Historical clinical and veterinary studies consistently describe lymph node palpation to detect enlargement as a key step in *T. brucei* diagnosis and follow-up (Boatin et al., 1986; Lutumba et al., 2005; Noireau et al., 1987; Nyimba et al., 2015; Vanhecke et al., 2010). Equally, lymph node aspirates allowing the isolation of *T. brucei* have been routinely performed for decades (Legros et al., 1999; Mumba Ngoyi et al., 2014; Scott et al., 1991). Around 100 studies published since the early 1900s have addressed different aspects of the interaction between *T. brucei* and the host lymphatic system in multiple mammals including humans, non-human primates, horses, dogs, donkeys, sheep, rabbits, cattle, and experimental animal models including rats and mice. In these studies, the lymph node-*T. brucei* interplay has seen many directions. One of the main directions, is the hematological and immunological consequences of host infection, particularly, immune population changes in lymphoid organs (and ensuing changes in the periphery) of infected hosts (Baker and Taylor, 1971; Barry and Emergy, 1984; Brown and Losos, 1977; Gasbarre et al., 1981, 1980; Hu, 1934; Ikede et al., 1977; B O Ikede and Losos, 1972; B. O. Ikede and Losos, 1972; Luckins, 1972; Magez et al., 2002, 1997; McCully and Neitz, 1971; Moulton and Sollod, 1976; M. Murray et al., 1974; P. K. Murray et al., 1974b, 1974a; Namangala et al., 2009, 2000b, 2000a; Okomo-Assoumou et al., 1995; Olsson et al., 1991; Sileghem et al., 1989; van den Ingh, 1977). This has led to a wealth of information on naive and adaptive immune responses to, and the function of many immune cell types during *T. brucei* infection. Other directions that have been explored under the lymph node-*T. brucei* umbrella include lymph node pathology (Barrowman and Roos, 1979; Morrison et al., 1981), lymph node invasion and its importance for tissue distribution (Alfituri et al., 2020a, 2020b; Rudin et al., 1984), and diverse variant antigen types present across the host body (Turner et al., 1986).

Despite this wealth of work, several questions remain unanswered with regards to the relevance of *T. brucei* presence in the lymphatic system in terms of overall relevance to host pathology (beyond alterations to immune system homeostasis); immune function and parasite recognition/elimination; and parasite heterogeneity. For the latter, it remains important to know whether the parasite population in the lymphatic system differs in any way from the parasite population in blood, and if it does, what the relevance of this difference is to infection and/or transmission. Between 5 and 4 decades ago, Ssenyonga and Adam, Hecker *et al*, and Tanner *et al* (Hecker and Brun, 1982; Ssenyonga and Adam, 1975; Tanner et al., 1980) did the first (and to our knowledge, the only) morphological analysis of *T. brucei* in the lymph nodes. They showed for the first time that parasites in the lymph nodes had a morphology that was intermediate between the conventional slender and stumpy forms found in blood. They also proposed lymph nodes as important sites for parasite replication and antigenic variation. Taking this work as a basis, in our work, our main aim was to investigate whether parasite heterogeneity exists between 21 lymph node sets in the mouse and whether (and how) it differs from the *T. brucei* population in blood. We considered this study as relevant given findings done over the last decade on parasite tropism and motility (reviewed in (Bargul et al., 2016; Crilly and Mugnier, 2021; Heddergott et al., 2012; Krüger et al., 2018; Krüger and Engstler, 2018; Silva Pereira et al., 2019), and the existence of modern tools allowed us to follow up the parasite populations with high spatial and temporal resolution (reviewed in De Niz et al., 2019), previously not available.

We used *in vivo* and *ex vivo* microscopy in murine models of infection to investigate 3 key questions namely 1) are *T. brucei* parasites equally distributed across all lymph nodes, or are some lymph nodes more enriched than others, and how does this compare to blood? 2) are *T. brucei* parasites different to those in blood in terms of stumpy/slender ratios, cell cycle progression, morphology and motility? 3) what is the impact of lymph node colonization on disease progression and, ultimately, mouse survival? We investigated all 3 questions at high-temporal resolution, given the characteristic cyclical pattern of infection, and reached conclusions that incorporate the time element as a key factor. We explore our findings in the context of each question below.

We found that parasite density in lymph nodes is different to that in blood both, in absolute numbers and considering temporal variations. The lymph node *T. brucei* population displays cyclical patterns that do not match those occurring in blood, showing a possible phase shift. We hypothesize that there is an interplay between the lymph node population and the blood population, with one affecting the other, although this remains a subject that demands further work in the future. While Ssenyonga and Adam suggested in 1975 (Ssenyonga and Adam, 1975) that a monomorphic trypanosome population is constantly flushed from the lymph to the blood, in our work we are unable to prove this hypothesis. However, the fact that a large live parasite population is present in the lymph nodes during parasitemia remission in the blood, and vice versa, suggests inter-organ communication that ensures parasite survival in separate anatomical locations. Our findings using LYVE-1-blocking antibodies, further support the possibility that the parasite population from either anatomical location is capable of replenishing the other. Also within the umbrella of this question, a further finding from our work is that parasite density and survival within different lymph node sets, is different. Namely, the lymph nodes located in the pelvis and abdomen are more heavily colonized than those in the limbs, head and neck, and thorax. Notably, our lab has previously mapped the largest extravascular reservoirs to organs in the pelvis and abdomen, i.e. the pancreas and gonadal adipose tissues (De Niz et al., 2021; Trindade et al., 2016). The abdominal and pelvic lymph nodes, together, help eliminate infections or inflammation in these regions, as well as drain inflammatory debris away from these regions to the systemic circulation. Further work could look into molecular, cellular and biophysical factors (eg. signaling, quorum sensing, and flows occurring between both organs, among others) mediating parasite migration between organs and lymph nodes.

Addressing our second question we found that the parasite population in the lymph nodes is significantly different to the one in blood. This is the case even considering major fluctuations through time of infection, observed across all factors analysed. Our findings are in agreement with previous reports (Hecker and Brun, 1982; Ssenyonga and Adam, 1975; Tanner et al., 1980), that defined the *T. brucei* lymph node population in the rat lymph nodes as a fast-replicating population with intermediate morphology. We here confirm that this is conserved in mice, and is consistent across lymph nodes. Nevertheless, considering all factors analysed (replication rate, stumpy/slender ratios, area, length, width, and speed) differences across lymph nodes in the 5 anatomical locations analysed exist, with most factors being similar across the abdominal and pelvic lymph nodes on one hand, and across the head and neck, limbs and thorax lymph nodes on the other hand. Strikingly, though, our findings have important implications for our understanding of *T. brucei* survival, morphology and behaviour, showing that these factors are heavily influenced by the microenvironment in which the parasites are present. This is the focus of further work in our lab addressing other extravascular reservoirs. While Tanner et al already raised in 1980 the question of whether only a specific parasite population is capable of invading organs and tissues beyond the blood, this question remains unanswered in our work, and will be an important venue of our research in the future. Even more so in the current context of research aiming to understand how the overall *T. brucei* population is built, and the implications thereof (Beaver et al., 2023; Briggs et al., 2021).

It remains to be understood why parasites in lymph nodes are generally bigger and faster than parasites in blood (despite major fluctuations with time possibly linked to parasite replication and influx/efflux across organs and blood). Previous work has suggested that faster swimmers are better able to eliminate host antibodies. Therefore, a possible hypothesis is that faster movement in lymph nodes is necessary for parasite survival. However, other alternatives exist including escape from immune cells, response to alternative signals present in the lymph nodes but not the blood, or the presence of exogenous components stimulating movement including the effect of forces acting on lymph flow. It is possible to hypothesize that both, host and parasite factors are involved in regulating parasite adaptation. Biophysical and biomechanical components in the two major tissues investigated in this work are largely different in terms of extracellular matrix, collagen presence, viscoelasticity, and the presence and nature of flow related to blood and lymphatic transport (Magno et al., 2015; O’Melia et al., 2019; Rohner et al., 2015; Wiig and Swartz, 2012). Other possible factors include those related to infection itself, such as changes in viscosity and temperature. Whether and how these parameters might influence parasite adaptation, remains an open question. Additional to this important point, another observation arising from this work worth following up was the relatively low proportion of stumpy forms found in the lymph nodes. Whether this arises because stumpy forms are incapable of invading lymph nodes, or because lymph nodes do not support differentiation, or stumpy form survival, this finding has important implications for our understanding of *T. brucei* transmission. So far, the field is most familiar with slender/stumpy fluctuations in the blood, but we know very little about these fluctuations at other organ and sub-organ levels. Our findings here add to the large wealth of work being generated to understand the relevance of tissue reservoirs for overall parasite survival and transmission potential.

Finally, in response to our third question, we found that both, lymph nodes and the lymphatic vasculature are highly remodeled throughout infection. While we have shown that remodeling of equivalent nature occurs in other organs (De Niz et al., 2021; Machado et al., 2022; Trindade et al., 2016), it is important to consider the implications of lymphatic system compromise. Although *T. brucei* is not considered *per se* a lymphatic disease as lymphatic filariasis is, it induces remarkable changes in systemic homeostasis as shown by the presence of major oedema (composed of heavily parasitized fluid) throughout the whole mouse body. This suggests alterations in lymphatic transport at a whole body level, which might have important implications for individual organ homeostasis (reviewed in (Stritt et al., 2021)) and overall immune function (reviewed in (Liao and Padera, 2013)), to name some factors. Altered lymphatic flow has been the subject of study in multiple other diseases (Cueni and Detmar, 2008; Liao and Padera, 2013; Mallick and Bodenham, 2003; Padera et al., 2016; Thomas et al., 2016) and is certainly an exciting new venue of research for *T. brucei*.

Altogether, we have found that lymph nodes harbour a unique population of *T. brucei* parasites which display significant fluctuations related to infection progression – suggesting adaptation processes similar to those occurring in the blood. Moreover, we show that while lymphatic system alterations have not been the focus of studies aiming to understand overall host pathology related to nagana or Human African trypanosomiasis, they might be key to our understanding of the big picture of host-pathogen interactions, immune response to *T. brucei,* and overall host pathology resulting from possible lymphadenopathy. Our work on lymph nodes is complementary to our studies of parasite heterogeneity in other extravascular sites, currently in progress. We are sure these findings pose an exciting basis for future research on *T. brucei* heterogeneity, parasite transmission, immunopathology and host-parasite biophysics and biomechanics.

## Materials and Methods

### Animals models

Animal experiments were performed according to EU regulations and approved by the Animal Ethics Committee of Instituto de Medicina Molecular (IMM) (AEC_2011_006_LF_TBrucei_IMM). Mice used across this study included wild-type C57BL/6J mice obtained from Charles River, France. Breedings were generated in-house at IMM. All mice were 6–9 weeks old males and females, with an average weight ranging between 25 and 30 g.

### Trypanosoma brucei parasites and infections

Mice were infected by intraperitoneal injection of either 3,000 *T. brucei* AnTat 1.1^E^ chimeric triple reporter parasites (Calvo-Alvarez et al., 2018) expressing the red-shifted firefly luciferase protein PpyREH9, TdTomato and Ty1 or GFP::PAD1-expressing *T. brucei* AnTat 1.1^E^. Based on our previous findings on the time of highest disease recrudescence in the survival experiments (De Niz et al., 2021), most experiments were capped to 20 days of infection, except those relevant to terminal body fluid and oedema measurements. Experiments relative to terminal body fluid were done with infections of 2,000 *T. brucei* EATRO1125 AnTat 1.1E 90-13 parasites.

### Intravital and ex vivo imaging

For intravital imaging, surgeries were performed as described in De Niz et al 2019a-c, De Niz et al 2020 for the various anatomical locations. Briefly, mice were anaesthetized with a mixture of ketamine (120 mg/kg) and xylazine (16 mg/kg) injected intraperitonially. Following verification of lack of reflex, mice were intraocularly injected with Hoechst 33342 (stock diluted in dH_2_O at 100 mg/ml; injection of 40 μg/kg mouse), 70 kDa FITC-Dextran (stock diluted in 1x PBS at stock concentration of 100 mg/ml; injection of 500 mg/kg), and LYVE-1 conjugated to A647 (clone 223322 R&D systems, used at 20 μg). A temporary glass window (Merk rectangular coverglass, 100 mm x 60 mm or circular coverglass (12 mm)) was implanted in each organ, and secured either surgically, with surgical glue, or via a vacuum, in order to enable visualization of the lymph node surface. All imaging was done in a Zeiss Cell Observer SD (spinning disc) confocal microscope (Carl Zeiss Microimaging, equipped with a Yokogawa CSU-X1 confocal scanner, an Evolve 512 EMCCD camera and a Hamamatsu ORCA-flash 4.0 VS camera) or in a 3i Marianas SDC (spinning disc confocal) microscopy (Intelligent Imaging Innovations, equipped with a Yokogawa CSU-X1 confocal scanner and a Photometrics Evolve 512 EMCCD camera). Laser units 405, 488, 561 and 640 were used to image Hoechst in nuclei, draining lymph nodes and lymphatics marked by FITC-Dextran and GFP::PAD1 in *T. brucei*, TdTomato in *T. brucei,* and LYVE1 in the lymphatic vascular endothelium, respectively. The objective used to image parasite density and vascular morphology was a 40x LD C-Apochoromat corrected, water immersion objective with 1.1 NA and 0.62 WD. The objective used to determine all cell cycle, survival, morphological and behavioural characteristics of *T. brucei* was a 100x plan-apochromat, oil immersion objective with 1.4 NA and 0.17 WD. Between 20 and 100 images were obtained in any one time lapse, with an acquisition rate of 20 frames per second. Since not all lymph nodes were easily accessible by intravital surgery, in order to gain access to all lymph node sets, we performed *ex vivo* imaging by excising the lymph node. For this, we performed z stacks consisting of 16 stacks covering up to 200 μm of tissue depth. For all acquisitions, the software used was ZEN blue edition v.2.6 (for the Zeiss Cell Observed SD) allowing export of images in .czi format, and 3i Slidebook reader v.6.0.22 (for the 3i Marianas SD), allowing export of images in TIFF format.

### Lymph node identification and measurement

Lymph nodes were identified as described by Van den Broeck *et al* (Van den Broeck et al., 2006) and Harrell *et al* (Harrell et al., 2008), by injection of subcutaneous, intrahepatic, and intradermal injection of Evans Blue dye. Result were confirmed by injection of 70 kDa FITC-Dextran whenever intravital microscopy was possible. The initial characterization was performed in infected mice at day 10 post-infection, when lymph nodes were enlarged, ensuring their correct identification from surrounding tissue. Confirmation of correct lymph node identification was performed by microscopy and comparison of anatomical location to the detailed mapping described by Van den Broeck (Van den Broeck et al., 2006).

#### Parasite density and diameter quantification

In order to quantify lymphatic vascular density for normalization of parasite load (i.e. expression as parasites per cm^2^ of vessel), we took as reference the vascular marker LYVE1 -A647. We segmented the area demarcated by LYVE.1, and quantified parasites within this area. We then expressed parasite number as a function of the area. Results shown in Tables 1 and 2 are the average of at least 100 measurements in separate fields of view. Vascular density measurements were performed throughout 20 days of infection. Vessel diameters and areas were measured using Fiji (Schneider et al., 2012). To identify the lymphatic vasculature, we intravenously injected 20 μg of antibodies against LYVE-1 (BioLegend) conjugated to A647, into infected mice at each day of infection, as previously described in the context of parasitology for CD31 (De Niz et al., 2021, 2018; Hopp et al., 2015)

### Quantification of parasite survival, cell cycle progression and stumpy/slender ratios

Parasite survival was defined by two features: absolute absence of parasite motility (not only lack of displacement but also lack of flagellar beating) and markers of cell death including blebbing or parasite destruction. Results shown in Table 3 are the percentage of live parasites in at least 100 fields of view and 3 independent mice, measured throughout 20 days of infection. Cell cycle progression was characterized by intravenous injection of Hoechst 33342 – this results in the labeling of the parasite nucleus (N) and kinetoplast (K), allowing for their quantification. Parasites were classified into 3 groups: 1 kinetoplast, 1 nucleus (1K1N), 2 kinetoplasts, 1 nucleus (2K1N), and 2 kinetoplasts, 2 nuclei (2K2N). Results shown in table 4 are the percentage of live parasites in at least 100 fields of view and 3 independent mice, measured throughout 20 days of infection. While all other experiments were performed using a TdTomato reporter line, stumpy/slender ratios were measured by the use of a GFP::PAD1 line, in GFP is expressed in parasites differentiating into stumpy forms or those fully differentiated. While we did not make this distinction in our work, we consider the classification as a proxy. For this reason, most results are expressed as proportion of true slender forms (i.e. those in which no GFP expression was detected and morphological characterization excluded a stumpy phenotype). Results shown in table 5 are the percentage of true slender forms (non-GFP-expressing and with non-stumpy morphology) in at least 100 fields of view and 3 independent mice, measured throughout 20 days of infection.

### Morphological and behavioural analysis

Morphological analyses were performed using Fiji (Schneider et al., 2012). Parasite width was calculated in the TdTomato reporter *T. brucei* line by measuring parasite diameter at the widest point (close to the location of the nucleus/nuclei). Parasite length was measured from the flagellar tip to the parasite anterior (from edge to edge). While in some cases the full area of the parasite was visible this was not always the case. In order to avoid confounding resulting from the parasite’s positioning, we calculated area for all parasites as the proxy of width by length. This will in all cases be slightly higher than the true area value, which varies based on the proportion of the free flagellum. Results shown in tables 6, 7 and 8 are the average area of at least 50 parasites in at least 3 independent mice measured throughout 20 days of infection. Behavioural analysis was limited to the measurement of parasite speed. Between 20 and 100 images were obtained in any one time lapse, with an acquisition rate of 20 frames per second. Speed was calculated as the function of distance over time. Results shown in table 9 are the average speed of at least 50 parasites in at least 3 independent mice measured throughout 20 days of infection.

### Quantification of oedema-related parasitemia

In infected mice, once oedemas began to form, free fluid could be easily acquired. At least 1µl of fluid was obtained from the abdominal-pelvic or thoracic region of the mice and diluted in 200 µl of 1x PBS. Parasite load was quantified using disposable hemocytometers. Results shown in table 14 are the average parasitemia in 4 independent mice measured throughout 20 days of infection.

### Measurements of percentage of body fluid mass

Free fluid mass was determined using a 6.2 MHz time-domain nuclear magnetic resonance small animal body composition analyzer (Minispec LF65, Bruker). Data were normalized to free fluid of mice prior to infection.

### Blocking lymphatic endothelial receptor LYVE-1

To investigate the effects of blocking LYVE-1, we used the antibodies MAB2125 (R&D systems, 20 μg per mouse), and Mouse IgG2 isotype controls were used as controls (eBioscience). Antibodies were injected intravenously every 2 days by tail vein injection, starting on the day of infection, and continuing until day 10 post-infection.

### Hierarchical clustering analysis

Hierarchical clustering analysis was performed using the Heatmapper tool developed by Babicki et al (Babicki et al., 2016)available at http://heatmapper.ca. Dendrograms and heatmaps were generated using the Expression tool. For all cases we used average linkage as the clustering method, and Euclidean distance measurement method. The distance and correlation matrices in Figure 4 were generated using the Pairwise tool, also based on Euclidean distance measurement.

### Value normalization and feature selection

Data normalization (shown in Table 10) was performed for all data shown in Tables 1 and 3-9 using the formula (x − 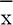)/S where x is the data value (the global average of each criteria), 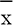 is the mean of the dataset (considering all organs analysed), and s is the standard deviation of the dataset. The normalized value shows how many standard deviations each datapoint is from the mean, and whether it’s greater (if the value is above 0) or less (if the value is below 0) than the mean. Feature selection was done by calculating the mean absolute deviation (MAD), which can be obtained by the formula *MAD* = *median* (|*x_i_*−*x_m_*|). *x_i_* is the i^th^ value in the dataset, while *x_m_* is the median value in the dataset. While feature selection can be used to discard features in a complex analysis, in our work we used this analysis to highlight the most significant factors for the distinction between LN and blood *T. brucei* populations.

### Quantification and statistical analysis

Data were displayed in graphs and heatmaps generated using Prism 9 software (GraphPad). Means, medians, survival, correlation tests, comparison tests, and error measures were calculated from triplicate experiments with 3 biological replicates each, and/or at least 100 images per condition. Means or geometric means were calculated for each section, as appropriate. For comparisons of measurements between LN and LN groups we performed multiple t tests in addition to a one-way ANOVA (differences were considered significant when p < 0.05). Pearson correlations measures (R), and R^2^ values were calculated to determine the strength of linear association between various parameters. For comparisons of survival, a log-rank (Mantel-Cox) test was performed, and p values < 0.05 were considered significant. Statistical details of experiments are included in the figure legends, the results section, and a Supplementary Table excel file. All data used for the generation of the figures is included as a supporting file.

## Acknowledgements

We thank Luisa Figueiredo and the Figueiredo lab at IMM for their helpful input and making this work possible. We thank Claudio Franco (IMM) and Virginie Uhlmann (EMBO) for their helpful input and advice on methods relevant to this work. We also thank Brice Rotureau (Institut Pasteur) for providing the triple reporter (TY1-TdTomato-FLuc) AnTat1.1^E^ *T. brucei* parasite line, and Fabien Guegan for generating the PAD1-GFP parasite line used in the lab. We thank Leonor Pinho for logistical help regarding experimental setups. We acknowledge the Rodent and Bioimaging Facilities of the Instituto de Medicina Molecular, especially the help of Jose Rino, Ana Nascimento and Antonio Temudo. This work was supported by HFSP (LT000047/2019-L) and EMBO (ALTF 1048-2016) to M.D.N. H.M. was funded by Fundação para a Ciência e Tecnologia (PD/BD/128286/2017). All work was carried out at the Figueiredo laboratory, funded by ERC (FatTryp, ref. 771714).

## Contributions

Conceptualization, M.D.N.; methodology, H.M., M.D.N.; software, M.D.N.; validation, H.M. and M.D.N.; formal analysis, H.M., M.D.N.; investigation H.M., M.D.N.; data curation, H.M., M.D.N.; writing – original draft, M.D.N; writing – review & editing, H.M., M.D.N.; visualization, M.D.N.; supervision, M.D.N..; project administration, M.D.N..; funding acquisition, H.M., M.D.N. (both members of Luisa Figueiredo’s lab).

## Supplementary Figure Legends

Figure S1. Analysis at high temporal resolution showed that during the first 24 hours after infection all lymph nodes are colonized faster than blood. **A)** 24 hours after infection they have, on average 8.2-fold (sub-cluster 1) and 25.5-fold (sub-cluster 2) higher parasite densities than blood (shown as a red dotted line). **B)** Parasite density in the blood rises quickly, rising by 192.5-fold by day 3 post-infection compared to day 1, while lymph nodes in sub-cluster 1 and sub-cluster 2 reach a relative increase of 30.1-fold and 25.4-fold respectively. **C)** The maximum parasite density across all days is reached in the second peak of parasitemia in the blood, but not in most lymph nodes. In most lymph nodes, the absolute maximum occurs during the first peak of parasitemia. Yet the difference between the maximum density at the first and second peaks of parasitemia is non-significant. **D)** The early stage of infection and the remission phase seem to favour lymph node invasion over blood, and throughout all 3 infection stages parasite density variations in the lymph nodes remain minimal. **E)** Although on average survival was highest in the blood, an absolute minimum of 12% survival was recorded in the blood, while the minimum survival in lymph node sub-clusters 1 and 2 was of 29.2 and 24.4% respectively. **F)** Survival fluctuations were highest in the blood compared to lymph nodes.

Figure S2. The lymph nodes are prohibitive for stumpy formation. A-B) During the first half of the infection, the average slender percentage in blood is 54.6%, while in the second half it is 21.3% reaching a minimum of 2%. In lymph node cluster 1 (HN and L), slender presence remains significantly higher throughout infection, with an average of 85.7% during the first 10 days, 74.9% during the last 10 days, and a minimum of 63% at any point during the infection. In lymph node cluster 2 (T, A and P), slender presence remains significantly higher than in blood throughout infection, with an average of 81.3% during the first 10 days, 67.4% during the last 10 days, and a minimum of 40% at any point during the infection (between days 16 and 20).

Figure S3. The *T. brucei* population in the lymph nodes is signifcantly longer and wider than the parasite population in the blood. **A)** Hierarchical clustering for width measurements showed the blood as a separate cluster to all lymph nodes, with no other significant clusters between the lymph node sets. **B)** Violin plots showing width distribution measurements for parasites in blood (red) and lymph nodes. **C)** Hierarchical clustering for length measurements showed the blood as a separate cluster to all lymph nodes, with no other significant clusters between the lymph node sets. **B)** Violin plots showing length distribution measurements for parasites in blood (red) and lymph nodes.

Figure S4. The *T. brucei* population in the lymph nodes is behaviourally different to the one in blood. Changes in speed were infection-time-dependent with drastic variations occurring within a short period of time during both parasitemia waves. **A)** During the first 10 days of infection, parasites in the blood had a median speed of 19.04 µm/s, while the lymph node parasite populations showed a median speed of 36.1 µm/s. **B)** During the last 10 days of infection, parasites in the blood had a median speed of 21.5 µm/s, while the lymph node parasite populations showed a median speed of 30.1 µm/s. **C-D)** Considering all time points, the maximum speed reached by parasites in any lymph node (50.92 µm/s), was significantly higher to the one reached in blood (27.46 µm/s).

## Notes

### Competing Interest Statement

The authors have declared no competing interest.

## Bibliography

Adamczyk LA, Gordon K, Kholová I, Meijer-Jorna LB, Telinius N, Gallagher PJ, van der Wal AC, Baandrup U. 2016. Lymph vessels: the forgotten second circulation in health and disease. Virchows Arch 469:3–17. doi:10.1007/s00428-016-1945-6

Alfituri OA, Bradford BM, Paxton E, Morrison LJ, Mabbott NA. 2020a. Influence of the Draining Lymph Nodes and Organized Lymphoid Tissue Microarchitecture on Susceptibility to Intradermal Trypanosoma brucei Infection. Front Immunol 11:1118. doi:10.3389/fimmu.2020.01118

Alfituri OA, Quintana JF, MacLeod A, Garside P, Benson RA, Brewer JM, Mabbott NA, Morrison LJ, Capewell P. 2020b. To the Skin and Beyond: The Immune Response to African Trypanosomes as They Enter and Exit the Vertebrate Host. Front Immunol 11:1250. doi:10.3389/fimmu.2020.01250

Babicki S, Arndt D, Marcu A, Liang Y, Grant JR, Maciejewski A, Wishart DS. 2016. Heatmapper: web-enabled heat mapping for all. Nucleic Acids Res 44:W147-53. doi:10.1093/nar/gkw419

Baker JR, Taylor AE. 1971. Experimental infections of the chimpanzee (Pan troglodytes) with Trypanosoma brucei brucei and Trypanosoma brucei rhodesiense. Ann Trop Med Parasitol 65:471–485. doi:10.1080/00034983.1971.11686780

Bargul JL, Jung J, McOdimba FA, Omogo CO, Adung’a VO, Krüger T, Masiga DK, Engstler M. 2016. Species-Specific Adaptations of Trypanosome Morphology and Motility to the Mammalian Host. PLoS Pathog 12:e1005448. doi:10.1371/journal.ppat.1005448

Barrowman PR, Roos JA. 1979. Lymph node pathology in Trypanosoma brucei-infected sheep. Onderstepoort J Vet Res 46:9–17.

Barry JD, Emergy DL. 1984. Parasite development and host responses during the establishment of Trypanosoma brucei infection transmitted by tsetse fly. Parasitology 88 ( **Pt 1**):67–84. doi:10.1017/s0031182000054354

Beaver AK, Crilly NP, Buenconsejo GY, Hakim JMC, Zhang B, Bobb B, Rijo-Ferreira F, Figueiredo LM, Mugnier MR. 2023. Extravascular spaces are the primary reservoir of antigenic diversity in Trypanosoma brucei infection. bioRxiv 2022.06.27.497797. doi:10.1101/2022.06.27.497797

Boatin BA, Wyatt GB, Wurapa FK, Bulsara MK. 1986. Use of symptoms and signs for diagnosis of Trypanosoma brucei rhodesiense trypanosomiasis by rural health personnel. Bull World Health Organ 64:389–395.

Briggs EM, Rojas F, McCulloch R, Matthews KR, Otto TD. 2021. Single-cell transcriptomic analysis of bloodstream Trypanosoma brucei reconstructs cell cycle progression and developmental quorum sensing. Nat Commun 12:5268. doi:10.1038/s41467-021-25607-2

Brown LA, Losos GJ. 1977. A comparative study of the responses of the thymus, spleen, lymph nodes and bone marrow of the albino rat to infection with Trypanosoma congolense and Trypanosoma brucei. Res Vet Sci 23:196–203.

Calvo-Alvarez E, Cren-Travaillé C, Crouzols A, Rotureau B. 2018. A new chimeric triple reporter fusion protein as a tool for in vitro and in vivo multimodal imaging to monitor the development of African trypanosomes and Leishmania parasites. Infect Genet Evol 63:391–403. doi:10.1016/j.meegid.2018.01.011

Capewell P, Cren-Travaillï¿½ C, Marchesi F, Johnston P, Clucas C, Benson RA, Gorman TA, Calvo-Alvarez E, Crouzols A, Jouvion G, Jamonneau V, Weir W, Lynn Stevenson M, O’Neill K, Cooper A, Swar NRK, Bucheton B, Ngoyi DM, Garside P, Rotureau B, MacLeod A. 2016. The skin is a significant but overlooked anatomical reservoir for vector-borne African trypanosomes. Elife 5:e17716. doi:10.7554/eLife.17716

Coles JA, Myburgh E, Ritchie R, Hamilton A, Rodgers J, Mottram JC, Barrett MP, Brewer JM. 2015. Intravital Imaging of a Massive Lymphocyte Response in the Cortical Dura of Mice after Peripheral Infection by Trypanosomes. PLoS Negl Trop Dis 9:e0003714. doi:10.1371/journal.pntd.0003714

Crilly NP, Mugnier MR. 2021. Thinking outside the blood: Perspectives on tissue-resident Trypanosoma brucei. PLoS Pathog 17:e1009866. doi:10.1371/journal.ppat.1009866

Cueni LN, Detmar M. 2008. The lymphatic system in health and disease. Lymphat Res Biol 6:109–122. doi:10.1089/lrb.2008.1008

De Niz M, Brás D, Ouarné M, Pedro M, Nascimento AM, Henao Misikova L, Franco CA, Figueiredo LM. 2021. Organotypic endothelial adhesion molecules are key for Trypanosoma brucei tropism and virulence. Cell Rep 36:109741. doi:10.1016/j.celrep.2021.109741

De Niz M, Meibalan E, Mejia P, Ma S, Brancucci NMB, Agop-Nersesian C, Mandt R, Ngotho P, Hughes KR, Waters AP, Huttenhower C, Mitchell JR, Martinelli R, Frischknecht F, Seydel KB, Taylor T, Milner D, Heussler VT, Marti M. 2018. Plasmodium gametocytes display homing and vascular transmigration in the host bone marrow. Sci Adv 4:eaat3775. doi:10.1126/sciadv.aat3775

De Niz M, Spadin F, Marti M, Stein J V, Frenz M, Frischknecht F. 2019. Toolbox for In Vivo Imaging of Host– Parasite Interactions at Multiple Scales. Trends Parasitol 35:193–212. doi:https://doi.org/10.1016/j.pt.2019.01.002

Gasbarre LC, Hug K, Louis J. 1981. Murine T lymphocyte specificity for African trypanosomes. II. Suppression of the T lymphocyte proliferative response to Trypanosoma brucei by systemic trypanosome infection. Clin Exp Immunol 45:165–172.

Gasbarre LC, Hug K, Louis JA. 1980. Murine T lymphocyte specificity for African trypanosomes. I. Induction of a T lymphocyte-dependent proliferative response to Trypanosoma brucei. Clin Exp Immunol 41:97–106.

Goodwin LG. 1970. The pathology of African Trypanosomiasis. Trans R Soc Trop Med Hyg 64:797–812. doi:10.1016/0035-9203(70)90096-9

Harrell MI, Iritani BM, Ruddell A. 2008. Lymph node mapping in the mouse. J Immunol Methods 332:170–174. doi:10.1016/j.jim.2007.11.012

Hecker H, Brun R. 1982. Comparative morphometric analysis of bloodstream and lymph forms of Trypanosoma (T.) brucei brucei grown in vitro and in vivo. Trans R Soc Trop Med Hyg 76:692–697. doi:10.1016/0035-9203(82)90241-3

Heddergott N, Krüger T, Babu SB, Wei A, Stellamanns E, Uppaluri S, Pfohl T, Stark H, Engstler M. 2012. Trypanosome motion represents an adaptation to the crowded environment of the vertebrate bloodstream. PLoS Pathog 8:e1003023. doi:10.1371/journal.ppat.1003023

Hopp CS, Chiou K, Ragheb DRT, Salman AM, Khan SM, Liu AJ, Sinnis P. 2015. Longitudinal analysis of Plasmodium sporozoite motility in the dermis reveals component of blood vessel recognition. Elife 4:e07789. doi:10.7554/eLife.07789

Hu CH. 1934. Studies on the Mature and Immature Lymphoid Cells of Spleen, Lymph Nodes and Thymus of Normal Rats and Rats Infected with Trypanosoma Brucei. Am J Pathol 10:29–42.5.

Ikede BO, Akpokodje JU, Hill DH, Ajidagba PO. 1977. Clinical, haematological and pathological studies in donkeys experimentally infected with Trypanosoma brucei. Trop Anim Health Prod 9:93–98. doi:10.1007/BF02236387

Ikede B. O., Losos GJ. 1972. Pathological changes in cattle infected with Trypanosoma brucei. Vet Pathol 9:272–277. doi:10.1177/030098587200900407

Ikede BO, Losos GJ. 1972. Spontaneous canine trypanosomiasis caused by T. brucei: meningo-encephalomyelitis with extravascular localization of trypanosomes in the brain. Bull Epizoot Dis Afr 20:221–228.

Jackson DG. 2019. Hyaluronan in the lymphatics: The key role of the hyaluronan receptor LYVE-1 in leucocyte trafficking. Matrix Biol 78–79:219–235. doi:10.1016/j.matbio.2018.02.001

Krüger T, Engstler M. 2018. The Fantastic Voyage of the Trypanosome: A Protean Micromachine Perfected during 500 Million Years of Engineering. Micromachines 9:63. doi:10.3390/mi9020063

Krüger T, Schuster S, Engstler M. 2018. Beyond Blood: African Trypanosomes on the Move. Trends Parasitol 34:1056–1067. doi:10.1016/j.pt.2018.08.002

Legros D, Evans S, Maiso F, Enyaru JC, Mbulamberi D. 1999. Risk factors for treatment failure after melarsoprol for Trypanosoma brucei gambiense trypanosomiasis in Uganda. Trans R Soc Trop Med Hyg 93:439–442. doi:10.1016/s0035-9203(99)90151-7

Liao S, Padera TP. 2013. Lymphatic function and immune regulation in health and disease. Lymphat Res Biol 11:136–143. doi:10.1089/lrb.2013.0012

Luckins AG. 1972. Adoptive immunity in experimental trypanosomiasis. Trans R Soc Trop Med Hyg 66:346–347. doi:10.1016/0035-9203(72)90231-3

Lutumba P, Robays J, Miaka C, Kande V, Simarro PP, Shaw APM, Dujardin B, Boelaert M. 2005. [The efficiency of different detection strategies of human African trypanosomiasis by T. b. gambiense]. Trop Med Int Health 10:347–356. doi:10.1111/j.1365-3156.2005.01391.x

Mabille D, Dirkx L, Thys S, Vermeersch M, Montenye D, Govaerts M, Hendrickx S, Takac P, Van Weyenbergh J, Pintelon I, Delputte P, Maes L, Pérez-Morga D, Timmermans J-P, Caljon G. 2022. Impact of pulmonary African trypanosomes on the immunology and function of the lung. Nat Commun 13:7083. doi:10.1038/s41467-022-34757-w

Machado H, Hofer P, Zechner R, Figueiredo LM. 2022. Adipocyte lipolysis protects the host against Trypanosoma brucei infection. bioRxiv 2022.11.05.515274. doi:10.1101/2022.11.05.515274

Magez S, Geuskens M, Beschin A, del Favero H, Verschueren H, Lucas R, Pays E, de Baetselier P. 1997. Specific uptake of tumor necrosis factor-alpha is involved in growth control of Trypanosoma brucei. J Cell Biol 137:715–727. doi:10.1083/jcb.137.3.715

Magez S, Stijlemans B, Caljon G, Eugster H-P, De Baetselier P. 2002. Control of experimental Trypanosoma brucei infections occurs independently of lymphotoxin-alpha induction. Infect Immun 70:1342–1351. doi:10.1128/IAI.70.3.1342-1351.2002

Magno R, Grieneisen VA, Marée AF. 2015. The biophysical nature of cells: potential cell behaviours revealed by analytical and computational studies of cell surface mechanics. BMC Biophys 8:8. doi:10.1186/s13628-015-0022-x

Mallick A, Bodenham AR. 2003. Disorders of the lymph circulation: their relevance to anaesthesia and intensive care. BJA Br J Anaesth 91:265–272. doi:10.1093/bja/aeg155

McCully RM, Neitz WO. 1971. Clinicopathological study on experimental Trypanosma brucei infections in horses. 2. Histopathological findings in the nervous system and other organs of treated and untreated horses reacting to nagana. Onderstepoort J Vet Res 38:141–175.

Morrison WI, Murray M, Sayer PD, Preston JM. 1981. The pathogenesis of experimentally induced Trypanosoma brucei infection in the dog. II. Change in the lymphoid organs. Am J Pathol 102:182–194.

Moulton JE, Sollod AE. 1976. Clinical, serologic, and pathologic changes in calves with experimentally induced Trypanosoma brucei infection. Am J Vet Res 37:791–802.

Mumba Ngoyi D, Ali Ekangu R, Mumvemba Kodi MF, Pyana PP, Balharbi F, Decq M, Kande Betu V, Van der Veken W, Sese C, Menten J, Büscher P, Lejon V. 2014. Performance of parasitological and molecular techniques for the diagnosis and surveillance of gambiense sleeping sickness. PLoS Negl Trop Dis 8:e2954. doi:10.1371/journal.pntd.0002954

Murray M, Murray PK, Jennings FW, Fisher EW, Urquhart GM. 1974. The pathology of Trypanosoma brucei infection in the rat. Res Vet Sci 16:77–84.

Murray PK, Jennings FW, Murray M, Urquhart GM. 1974a. The nature of immunosuppression in Trypanosoma brucei infections in mice. II. The role of the T and B lymphocytes. Immunology 27:825–840.

Murray PK, Jennings FW, Murray M, Urquhart GM. 1974b. The nature of immunosuppression in Trypanosoma brucei infections in mice. I. The role of the macrophage. Immunology 27:815–824.

Myburgh E, Coles JA, Ritchie R, Kennedy PGE, McLatchie AP, Rodgers J, Taylor MC, Barrett MP, Brewer JM, Mottram JC. 2013. In Vivo Imaging of Trypanosome-Brain Interactions and Development of a Rapid Screening Test for Drugs against CNS Stage Trypanosomiasis. PLoS Negl Trop Dis 7:e2384.

Namangala B, Brys L, Magez S, De Baetselier P, Beschin A. 2000a. Trypanosoma brucei brucei infection impairs MHC class II antigen presentation capacity of macrophages. Parasite Immunol 22:361–370. doi:10.1046/j.1365-3024.2000.00314.x

Namangala B, DE Baetselier P, Beschin A. 2009. Quantitative differences in immune responses in mouse strains that differ in their susceptibility to Trypanosoma brucei brucei infection. J Vet Med Sci 71:951–956. doi:10.1292/jvms.71.951

Namangala B, de Baetselier P, Brijs L, Stijlemans B, Noël W, Pays E, Carrington M, Beschin A. 2000b. Attenuation of Trypanosoma brucei is associated with reduced immunosuppression and concomitant production of Th2 lymphokines. J Infect Dis 181:1110–1120. doi:10.1086/315322

Noireau F, Gouteux JP, Duteurtre JP. 1987. [Diagnostic value of a card agglutination test (Testryp CATT) in the mass screening of human trypanosomiasis in the Congo]. Bull Soc Pathol Exot Filiales 80:797–803.

Nyimba PH, Komba EVG, Sugimoto C, Namangala B. 2015. Prevalence and species distribution of caprine trypanosomosis in Sinazongwe and Kalomo districts of Zambia. Vet Parasitol 210:125–130. doi:10.1016/j.vetpar.2015.04.003

O’Melia MJ, Lund AW, Thomas SN. 2019. The Biophysics of Lymphatic Transport: Engineering Tools and Immunological Consequences. iScience 22:28–43. doi:10.1016/j.isci.2019.11.005

Okomo-Assoumou MC, Daulouede S, Lemesre JL, N’Zila-Mouanda A, Vincendeau P. 1995. Correlation of high serum levels of tumor necrosis factor-alpha with disease severity in human African trypanosomiasis. Am J Trop Med Hyg 53:539–543. doi:10.4269/ajtmh.1995.53.539

Olsson T, Bakhiet M, Edlund C, Höjeberg B, Van der Meide PH, Kristensson K. 1991. Bidirectional activating signals between Trypanosoma brucei and CD8+ T cells: a trypanosome-released factor triggers interferon-gamma production that stimulates parasite growth. Eur J Immunol 21:2447–2454. doi:10.1002/eji.1830211022

Padera TP, Meijer EFJ, Munn LL. 2016. The Lymphatic System in Disease Processes and Cancer Progression. Annu Rev Biomed Eng 18:125–158. doi:10.1146/annurev-bioeng-112315-031200

Rohner NA, McClain J, Tuell SL, Warner A, Smith B, Yun Y, Mohan A, Sushnitha M, Thomas SN. 2015. Lymph node biophysical remodeling is associated with melanoma lymphatic drainage. FASEB J Off Publ Fed Am Soc Exp Biol 29:4512–4522. doi:10.1096/fj.15-274761

Rudin W, Pongponratn E, Jenni L. 1984. Electron-microscopic localization of Trypanosoma brucei gambiense transmitted by Glossina morsitans centralis in Microtus montane. Acta Trop 41:325–334.

Schneider CA, Rasband WS, Eliceiri KW. 2012. NIH Image to ImageJ: 25 years of image analysis. Nat Methods 9:671–675. doi:10.1038/nmeth.2089

Schuberg, A., Boeing W. 1913. Ueber den Weg der Infektion bei Trypanosomen - und Spirochätenerkrankungen. Dtsch Medizinische Wochenschrift 19:877–879.

Scott JA, Davidson RN, Moody AH, Bryceson AD. 1991. Diagnosing multiple parasitic infections: trypanosomiasis, loiasis and schistosomiasis in a single case. Scand J Infect Dis 23:777–780. doi:10.3109/00365549109024307

Sileghem M, Darji A, Remels L, Hamers R, De Baetselier P. 1989. Different mechanisms account for the suppression of interleukin 2 production and the suppression of interleukin 2 receptor expression in Trypanosoma brucei-infected mice. Eur J Immunol 19:119–124. doi:10.1002/eji.1830190119

Silva Pereira S, Trindade S, De Niz M, Figueiredo LM. 2019. Tissue tropism in parasitic diseases. Open Biol 9:190036. doi:10.1098/rsob.190036

Ssenyonga GS, Adam KM. 1975. The number and morphology of trypanosomes in the blood and lymph of rats infected with Trypanosoma brucei and T. congolense. Parasitology 70:255–261. doi:10.1017/s0031182000049714

Stritt S, Koltowska K, Mäkinen T. 2021. Homeostatic maintenance of the lymphatic vasculature. Trends Mol Med 27:955–970. doi:10.1016/j.molmed.2021.07.003

Tanner M, Jenni L, Hecker H, Brun R. 1980. Characterization of Trypanosoma brucei isolated from lymph nodes of rats. Parasitology 80:383–391. doi:10.1017/s0031182000000834

Thomas SN, Rohner NA, Edwards EE. 2016. Implications of Lymphatic Transport to Lymph Nodes in Immunity and Immunotherapy. Annu Rev Biomed Eng 18:207–233. doi:10.1146/annurev-bioeng-101515-014413

Trindade S, Niz M De, Sequeira M, Bizarra-Rebelo T, Bento F, Dejung M, Ferreira J, Butter F, Bringaud F, Gjini E, Figueiredo L. 2021. Persistence behavior in African trypanosomes during adipose tissue colonization. Nat Portf. doi:10.21203/rs.3.rs-499897/v1

Trindade S, Rijo-Ferreira F, Carvalho T, Pinto-Neves D, Guegan F, Aresta-Branco F, Bento F, Young SA, Pinto A, Van Den Abbeele J, Ribeiro RM, Dias S, Smith TK, Figueiredo LM. 2016. Trypanosoma brucei Parasites Occupy and Functionally Adapt to the Adipose Tissue in Mice. Cell Host Microbe 19:837–848. doi:https://doi.org/10.1016/j.chom.2016.05.002

Turner CM, Hunter CA, Barry JD, Vickerman K. 1986. Similarity in variable antigen type composition of Trypanosoma brucei rhodesiense populations in different sites within the mouse host. Trans R Soc Trop Med Hyg 80:824–830. doi:10.1016/0035-9203(86)90395-0

Van den Broeck W, Derore A, Simoens P. 2006. Anatomy and nomenclature of murine lymph nodes: Descriptive study and nomenclatory standardization in BALB/cAnNCrl mice. J Immunol Methods 312:12–19. doi:https://doi.org/10.1016/j.jim.2006.01.022

van den Ingh TS. 1977. Pathomorphological changes in Trypanosoma brucei brucei infection in the rabbit. Zentralblatt fur Vet R B J Vet Med Ser B 24:773–786. doi:10.1111/j.1439-0450.1977.tb00970.x

Vanhecke C, Guevart E, Ezzedine K, Receveur M-C, Jamonneau V, Bucheton B, Camara M, Vincendeau P, Malvy D. 2010. [Human African trypanosomiasis in mangrove epidemiologic area. Presentation, diagnosis and treatment in Guinea, 2005-2007]. Pathol Biol (Paris) 58:110–116. doi:10.1016/j.patbio.2009.07.033

Wiig H, Swartz MA. 2012. Interstitial fluid and lymph formation and transport: physiological regulation and roles in inflammation and cancer. Physiol Rev 92:1005–1060. doi:10.1152/physrev.00037.2011

