## Supplementary figures and images for "The lymphatic system favours survival of a unique *T. brucei* population, and its invasion results in major host pathology"

### Figure S1

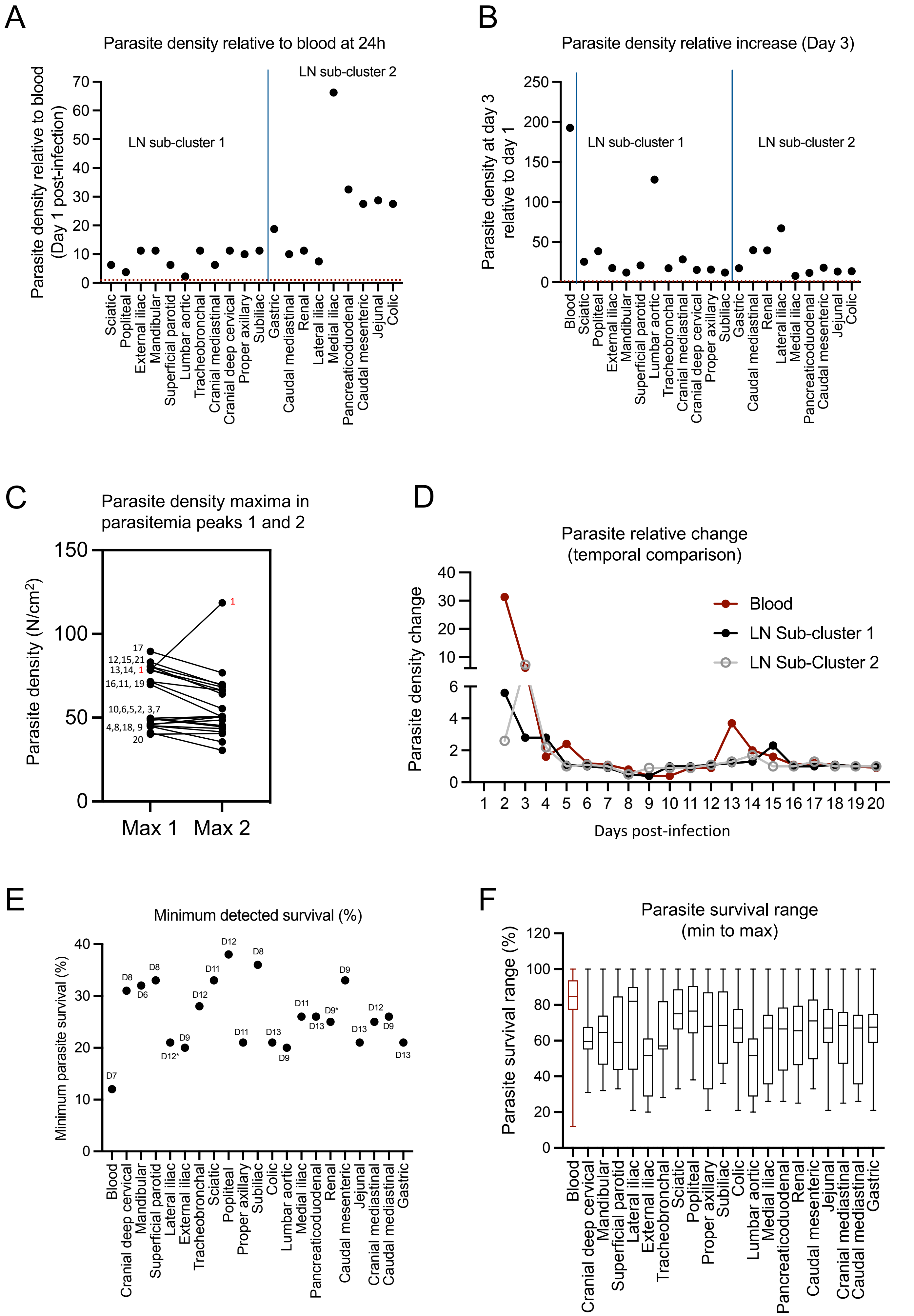

### Figure S2

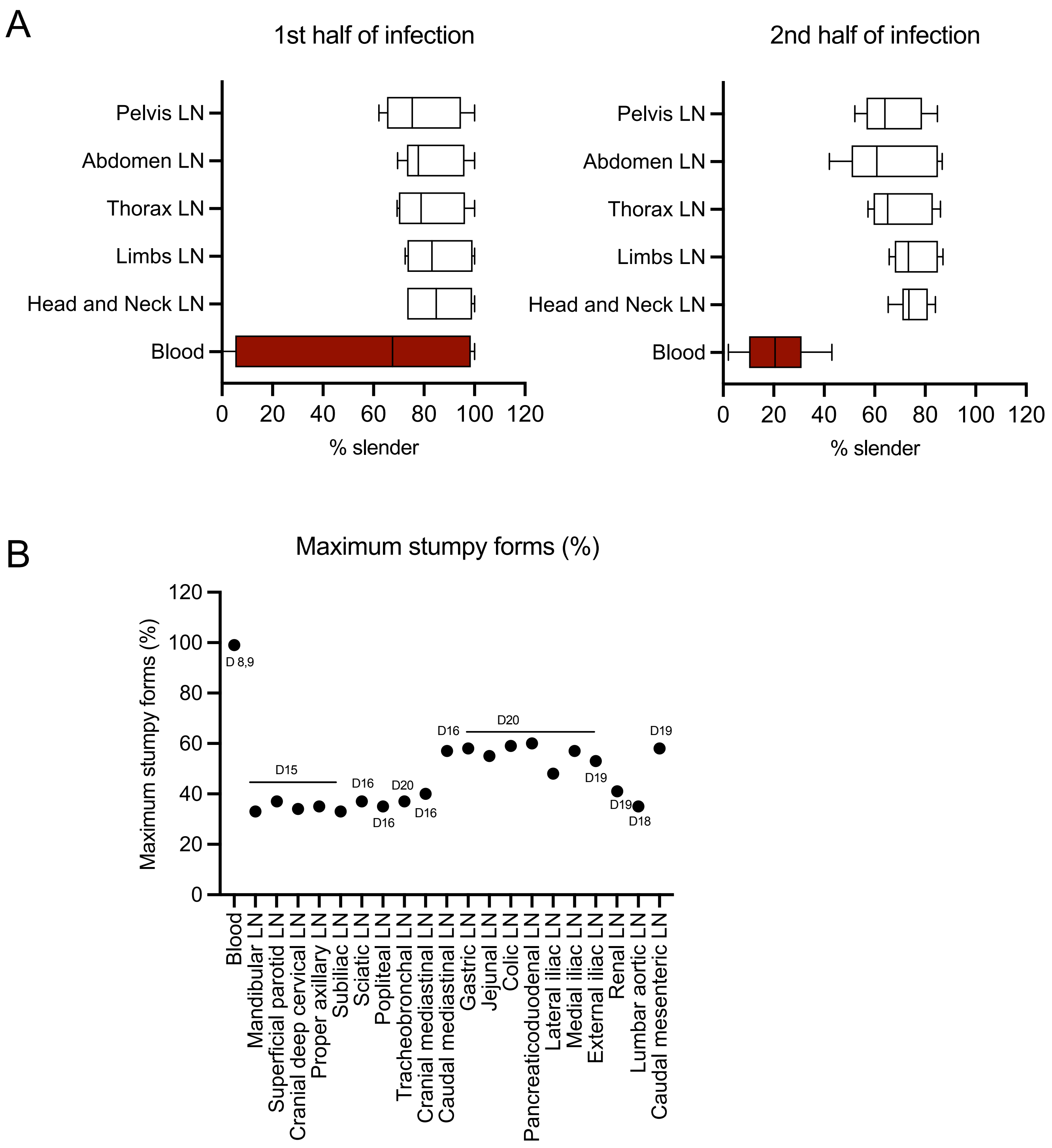

### Figure S3

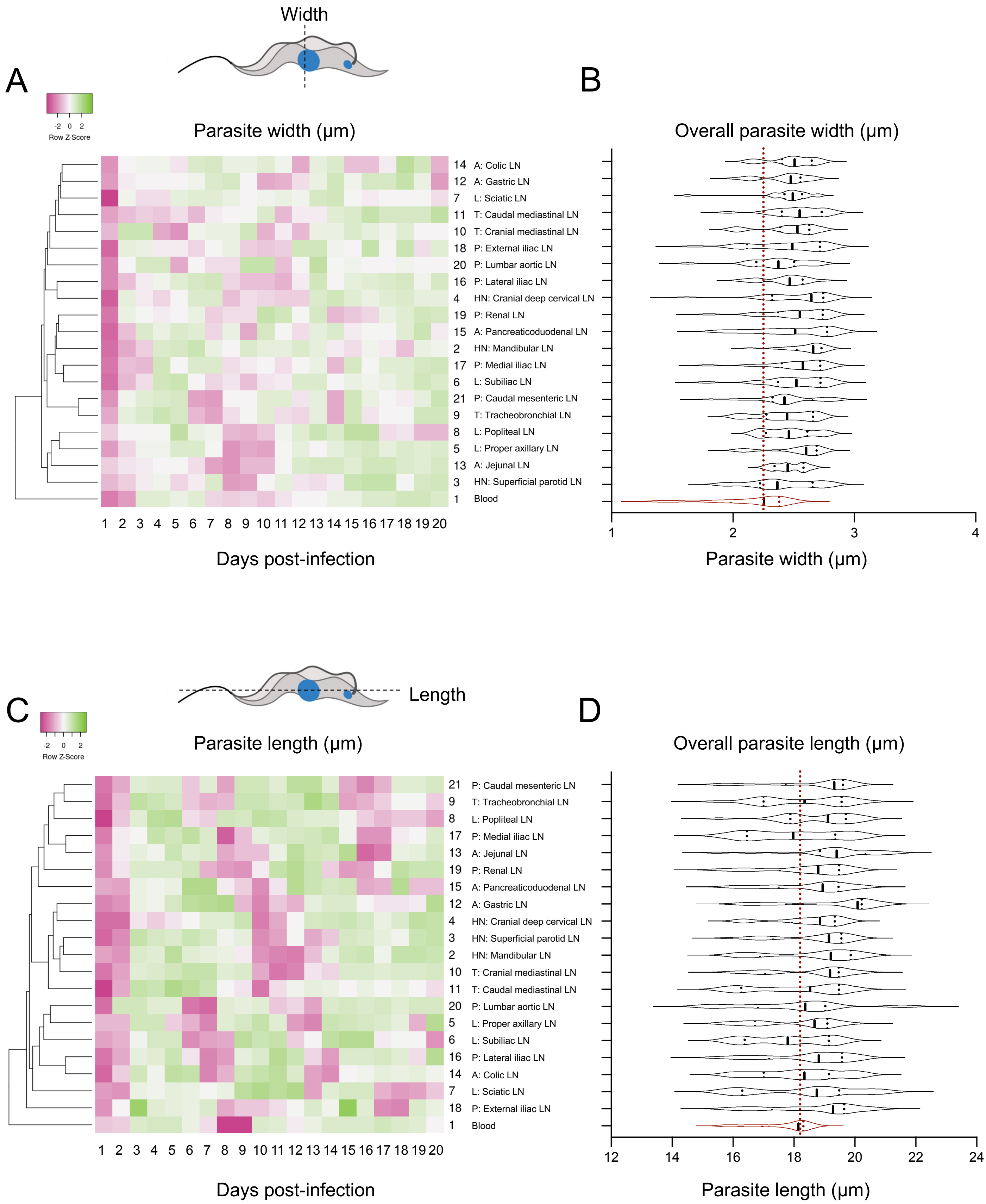

### Figure S4

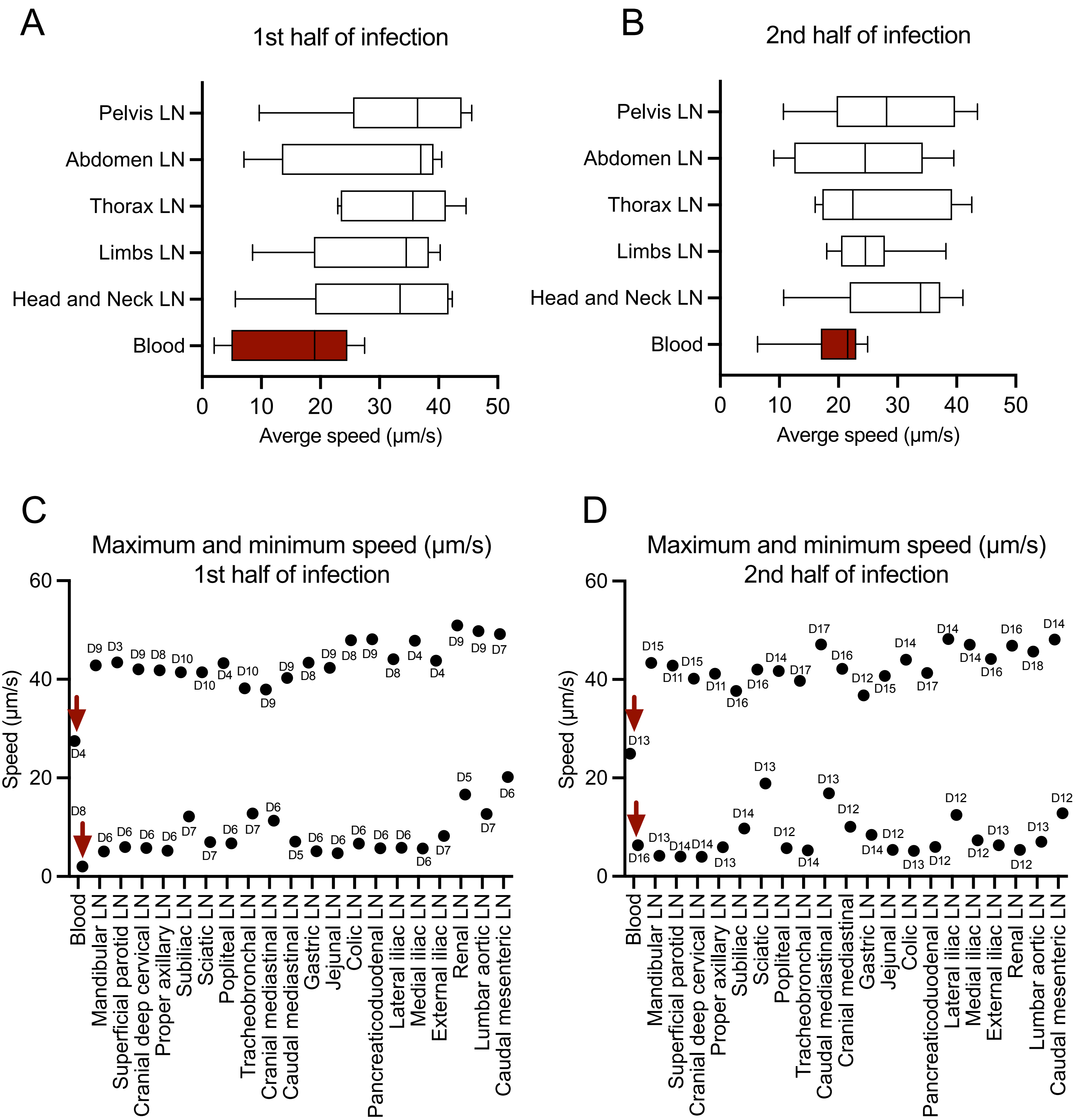
